# A framework for reconstructing ancient food webs using functional trait data

**DOI:** 10.1101/2024.01.30.578036

**Authors:** Jack O. Shaw, Alexander M. Dunhill, Andrew P. Beckerman, Jennifer A. Dunne, Pincelli M. Hull

## Abstract

1. Food webs provide quantitative insights into the structure and dynamics of ecological communities. Previous work has shown their utility in understanding community responses to modern and ancient perturbations, including anthropogenic change and mass extinctions. However, few ancient food webs have been reconstructed due to difficulties assessing trophic interactions amongst extinct species derived from an incomplete fossil record.
2. We present and assess the Paleo Food web Inference Model (PFIM). PFIM uses functional trait data—predictive of interactions in modern ecosystems and commonly available for fossil organisms—to reconstruct ancient food webs. We test the model by (i) applying it to four modern ecosystems with empirical constrained food webs to directly compare PFIM-constructed networks to their empirical counterparts, (ii) by carefully comparing discrepancies between PFIM-inferred and empirical webs in one of those systems, and (iii) by comparing networks describing feasible trophic interactions (“feasible webs”) with networks to which we superimpose characteristic interaction distributions derived from modern theory (“realized webs”). As a proof of concept, we then apply the method to faunal data from two Cambrian fossil deposits to reconstruct ancient trophic systems.
3. PFIM-inferred feasible food webs successfully predict ∼70% of trophic interactions across four modern systems. Furthermore, inferred food webs with enforced interaction distributions (i.e., realized webs) accurately predict ∼90% of interactions. Comparisons with a global database of trophic interactions and other food web models, suggest that under sampling of empirical webs accounts for up to 21% of the remaining differences between PFIM and empirical food webs.
4. Food webs can be reasonably approximated by inferring trophic interactions based upon life habit traits. This study provides the foundation to use trait-based inference models across the fossil record to examine ancient food webs and community evolution.

## INTRODUCTION

Ecological communities are complex systems composed of interacting organisms, and food webs are networks describing “who eats whom” in these systems. Such trophic networks—in which organisms are represented by nodes and trophic interactions are represented by directed links— have proven vital for disentangling the myriad of organismal- to community-scale ecological processes occurring in ecosystems (e.g., (Cardinale et al. 2009; Thompson et al. 2012; Albouy et al. 2019)).

Analyses of ancient food webs, considering hypothetical interactions between fossil taxa in an assemblage, have elucidated deep-time responses to ecological innovations (e.g., the proliferation of biomineralizing taxa across the Cambrian Explosion; (Dunne et al. 2008)), invasions (e.g., (Kempf et al. 2020)), and mass extinctions (e.g., (Roopnarine 2009; Dunne et al. 2014; Roopnarine and Dineen 2018)). Compared to typical analyses of fossil diversity, the food web perspective considers how interactions between organisms relate to dynamical properties of communities, which are otherwise unclear. Importantly, the geological literature contains numerous case studies of biotic responses to biotic and abiotic changes, such as “ecological avalanches” generated by primary extinctions (Vermeij 2004) and global warming (Mayhew et al. 2008), pertinent to the ongoing climate and biodiversity crises.

However, ancient food web analysis is relatively rare, due to difficulties and assumptions made in their construction, precluding testing of hypotheses about long-term community evolution.

Previous reconstructions of ancient food webs have generally taken one of two approaches to determine interactions between individual species while accounting for uncertainty generated by fossil record biases: (i) manually assigning interactions based on fossil evidence, modern analogs, phylogenetic conservation, and/or morphology (e.g., (Dunne et al. 2008, 2014)); and (ii) inferring interactions probabilistically between pre-defined trophic guilds (e.g., (Roopnarine 2006; Roopnarine and Angielczyk 2015; Kempf et al. 2020)). The first approach has the advantage of allowing for entirely novel network structures if the data suggest them, but it is generally labor intensive, data limited, and subjective. Although there are some instances of direct consumption evidenced by fossils (e.g., (Klug et al. 2021)), many inferences of interactions are based on evidence such as predation marks (e.g., bore holes and bite marks; (Klompmaker et al. 2017)) and gut contents (e.g., (Vannier 2012)). In most cases, however, interactions are commonly hypothesized based on modern analogs, phylogenetic conservation of trophic relationships, and functional traits evidenced in morphology (e.g., (Dunne et al. 2008)). The second approach, probabilistically distributing interactions, although less impacted by these disadvantages, is biased by assuming conserved community structure (i.e., the link distribution) between ancient and modern systems.

Predicting trophic interactions is difficult even in modern, information-rich assemblages (Poisot et al. 2015), and even more so in fossil assemblages, given the limited information preserved (e.g., marine fossil assemblages may capture a maximum ∼38% of their original diversity in the best-preserved environments (Shaw et al. 2021*a*)). A realized interaction between two taxa at one point in space or time may be absent or simply not observed at another point (Poisot et al. 2012). The realization of an interaction between two taxa depends on (i) sufficient local abundance such that individuals can interact (Petchey et al. 2008), (ii) trait matching between individuals such that one taxon can physically consume the other (Portalier et al. 2019), (iii) indirect interactions (including non-trophic interactions; (Trussell et al. 2017)) and (iv) environmental conditions (e.g., interactions between temperature and body-size; (Angilletta et al. 2004)).

There are several modern tools available to infer species interactions, although most are inapplicable to fossil data. For example, interactions may be inferred based on those observed elsewhere (e.g., (Pichler et al. 2020)) or DNA analyses of gut contents (e.g., (Roslin and Majaneva 2016)), methods that are not appropriate for ancient food webs. More mechanistic approaches link optimal foraging biology to body size to predict network structure (e.g., (Petchey et al. 2008; Portalier et al. 2019)). Although these approaches may be valuable, they assume that community structure and foraging behavior are preserved through deep time. A more neutral and feasible approach relies on a multivariate view of species functional traits. This type of data is available for fossil taxa and has the potential to explain variation in contemporary food web structures (Eklöf et al. 2013). Functional trait-based analyses are already commonly applied to the fossil record and have shown, for instance, increases in functional diversity between the Cambrian and modern (Bush and Bambach 2011) and correlations between extinction selectivity and functional diversity change (e.g., (Dunhill et al. 2018)).

For aquatic organisms, three life habit categories (motility, habitat depth relative to the substrate, and trophic mode) are commonly used to classify them into ecological modes of life (Bambach et al. 2007). While coarse, this approach reduces issues associated with finer delineations of traits, particularly when considering extinct organisms. As in modern assemblages (e.g., (Eklöf et al. 2013)), functional trait data have been used to consider ancient food web structure (e.g., (Kempf et al. 2020)), on the basis that functional ecology is connected to trophic interactions and food web structure (e.g., (Eklöf et al. 2013; Cirtwill and Eklöf 2018)).

Here we present and test the Paleo Food web Inference Model (“PFIM”; Fig. 1), a trophic interaction inference model utilizing taxon-level trait information (motility, tiering, feeding, and size). This framework builds upon previous work in which feasible trophic interactions are assumed between broad ecological categories, centered around “Bambachian megaguilds” (Roopnarine 2006), and species-level networks are generated through stochastic distribution of links among these groups (based on pre-defined structures). PFIM represents three notable advances on current inference-based models for paleontological applications. First, PFIM generates “feasible food webs”, reflecting all potential interactions that might occur over time and space according to a set of interaction rules. Feasible food webs are readily comparable to one another, permitting spatial and temporal evaluations of deep-time trophic trends, without any a priori assumptions about network structure. Second, PFIM provides a flexible framework for generating hypothetical “realized food webs” with varying network characteristics, such as link-species distributions, permitting analyses of the impacts of changes in food web structure on the dynamical properties of the system. PFIM builds on previous studies ((Roopnarine and Dineen 2018) and references therein), which distributed links amongst taxa based on the distribution of trophic interactions in modern food webs, to facilitate the comparison of different network structures in ancient communities. Third, PFIM is written in R, a commonly used coding language in ecology and paleontology, and is simple to execute.

**Figure 1:**
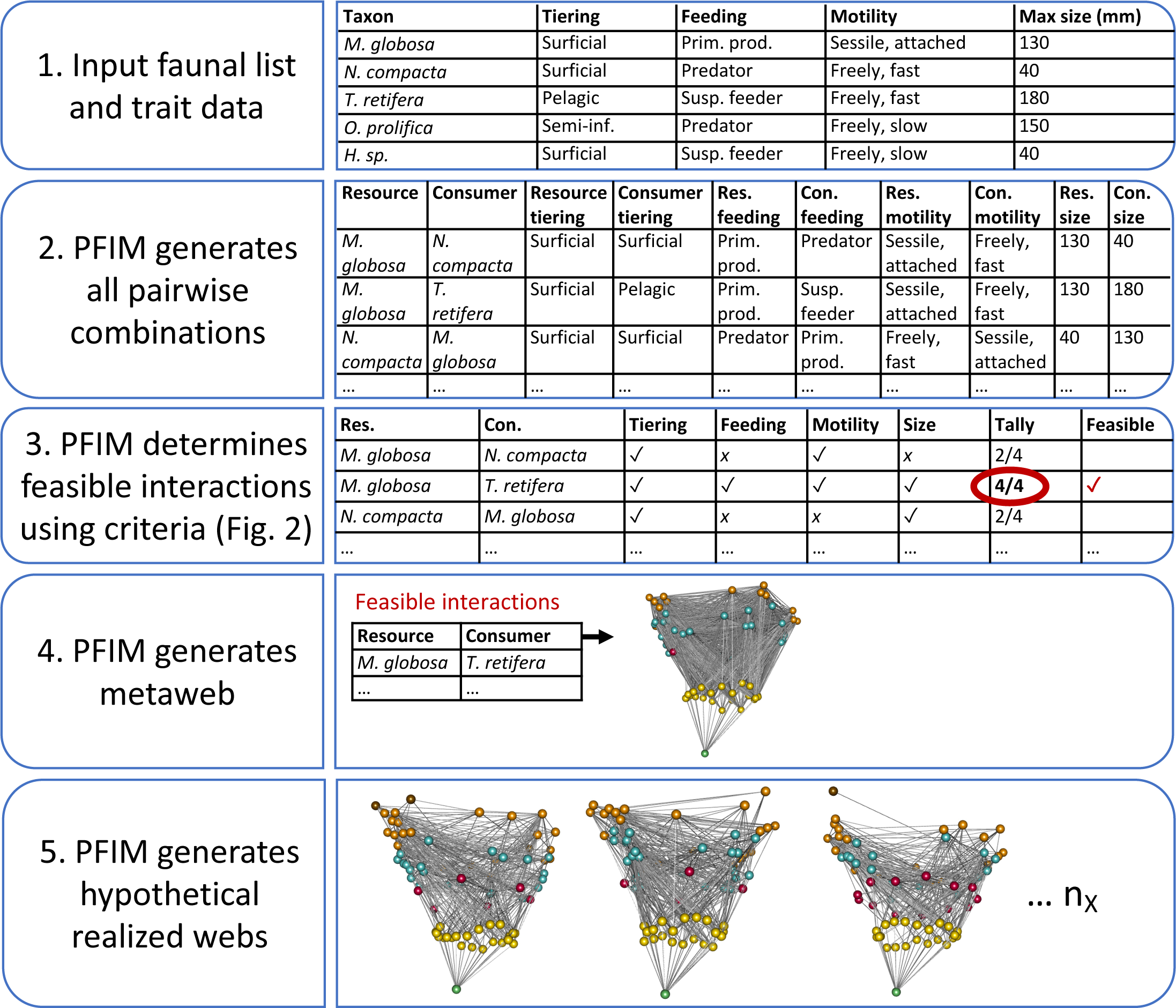
Schematic describing how PFIM reconstructs food webs, including feasible and realized webs, from faunal lists and functional traits.

In addition to presenting the model, we test its predictive accuracy by comparing PFIM- constructed networks to known modern food webs. We further investigate the predictive accuracy of PFIM by (i) analyzing whether our criteria for feasible interactions are sound and (ii) comparing our method to an existing inference method based on modern ecological data and theory. Furthermore, we apply PFIM to faunal data from two well-known Cambrian (∼500 Ma) deposits to explore differences between reconstructed ancient and modern food webs.

The supplementary information includes URLs linking to GitHub repositories containing (i) R code for PFIM and (ii) data and R code used for analyses in this paper.

## METHODS

### Model description

PFIM uses (i) motility, tiering, feeding, and size data to define a plausible food web based on encounter feasibility (e.g., describing all metazoan taxa that could feasibly interact based on their traits) and (ii) hypothesized link distributions, defined by ecological theory, to create a series of hypothetical realized webs.

#### Step 1: Construct feasible food web

Life habit (motility, tiering and feeding) and size—together referred to as functional trait data— are key predictors of consumer-resource interactions (e.g., (Eklöf et al. 2013; Gravel et al. 2013; Cirtwill and Eklöf 2018)) and are easily determined for fossil metazoans. We defined the feasibility of interactions between two taxa based on traits, creating a default set of interaction criteria (Figs. 1,2). For example, in terms of motility compatibility, “nonmotile, attached” taxa are considered unable to consume “motile, fast” taxa, but the reverse is feasible. Interaction criteria are consistent with several examples of established foraging theory (including consumer-resource size ratios), modern patterns of encounter probability (i.e., tiering), and functional constraints (i.e., motility and feeding) (e.g., (Beckerman et al. 2006; Petchey et al. 2008;

**Figure 2:**
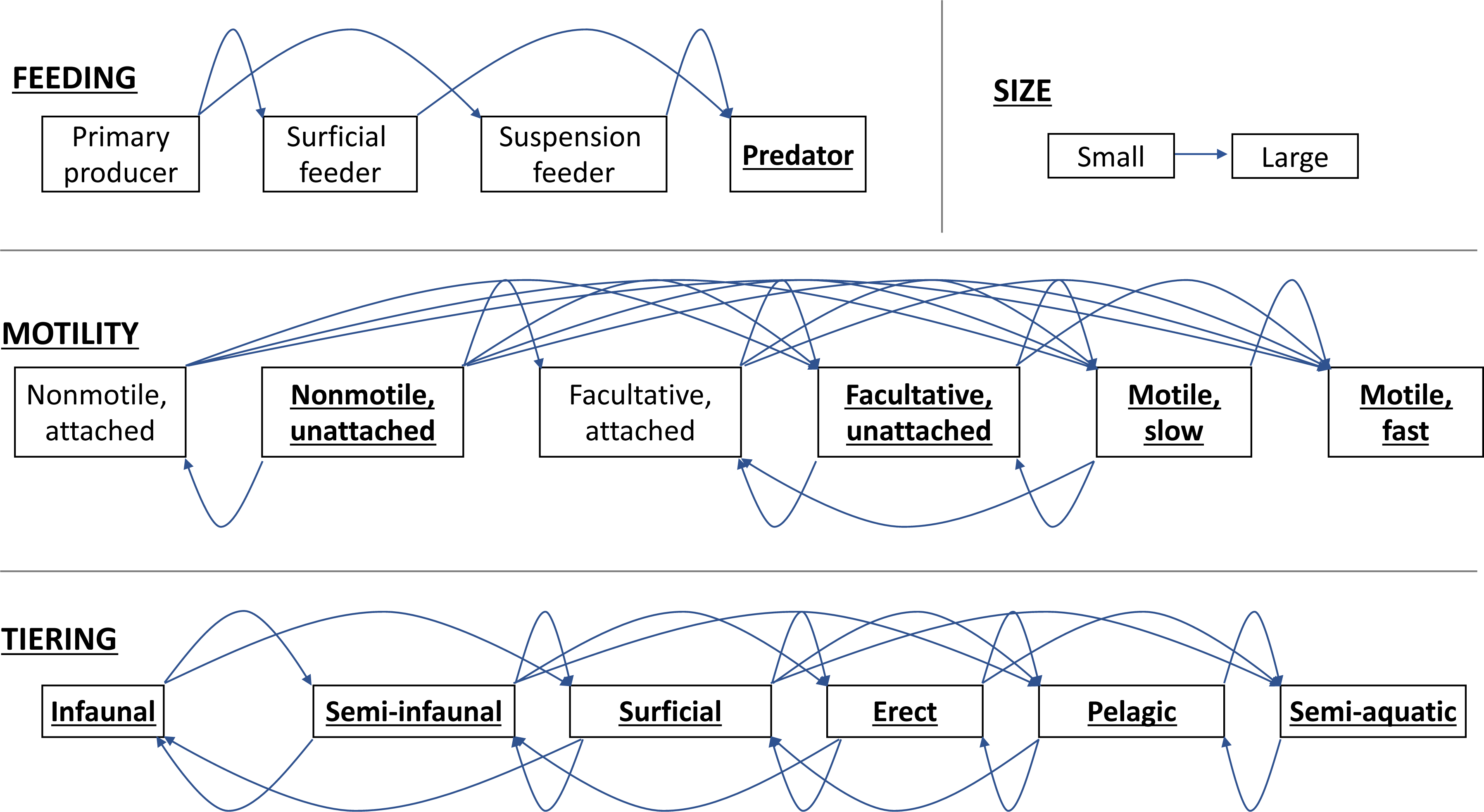
Interaction criteria used to determine if interactions are feasible between taxa belonging to different functional trait modes. Arrows point from resource to consumer. All lines above labels have arrows pointing to the right and all lines below the labels have arrows pointing to the left. Traits permitting cannibalism are underlined.

Portalier et al. 2019)). PFIM allows the user to replace the default interaction criteria set with custom, user-defined interaction criteria as needed.

The feasible food web inference process is as follows:

i. PFIM requires a data-frame listing metazoan taxon names as well as motility, tiering, feeding, and size. PFIM adds a basal node to represent primary producers (see supplementary information for more details), given that non-metazoans are commonly insufficiently recorded in metazoan fossil surveys.
ii. PFIM generates a pairwise list of all possible combinations of taxa in the faunal list and corresponding functional trait data.
iii. **Links**: PFIM applies the four interaction criteria (Fig. 2) to all pairwise combinations and adds a link between nodes when all four interaction criteria are met (see supplement for analyses of alternative criteria).
iv. **Feasible food web**: the network of all physically feasible links.

#### Step 2: Generate hypothetical realized webs

PFIM generates a series of replicate hypothetical realized webs using a link distribution function to reduce the feasible links assigned to each node to match a hypothetical distribution and number of realized links. The default link distribution function is a mixed exponential-power law in-degree distribution as proposed by (Roopnarine 2006) and used in numerous studies since (e.g., (Roopnarine and Dineen 2018)), although the user can define a custom distribution function. The selected distribution takes the form of *P*(*r*) = *e*^*r/E*^ where *r* is the in-degree of the consumer, 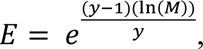, *M* is the total number of nodes, and *y* = 2.5. 1000 alternative hypothetical realized webs are generated by default (see (Roopnarine 2006) for an in-depth discussion of the rationale behind this equation). For the remainder of this paper we simply use the term “realized web” to refer to PFIM-inferred networks in which a mixed exponential-power law in-degree distribution is applied.

### Data compilation

We assessed the efficacy and utility of the PFIM by reconstructing four well-studied modern food webs and two ancient food webs using traits, interaction criteria, and link distributions (Table 1). These six webs were selected due to their broad higher-rank taxonomic compositions spanning multiple metazoan phyla (all webs contain annelids, arthropods, chordates, and mollusks; the number of phyla represented in each web ranges 4-9), and well-resolved taxonomic compositions (nodes resolved to family-level or below, with most taxa resolved to genus or species). We focused on aquatic assemblages given their prominence in the fossil record and the quality of available interaction data. Our comparisons include only four previously published modern localities, as many published modern food webs do not meet the criteria above (i.e., broad taxonomic compositions, high taxonomic resolution, and aquatic). Faunal and trophic interaction data for the two ancient localities—the Burgess and Chengjiang Shales—were taken from a previous publication that manually designated pairwise interactions between taxa based on expert opinion (Dunne et al. 2008).

**Table 1:**
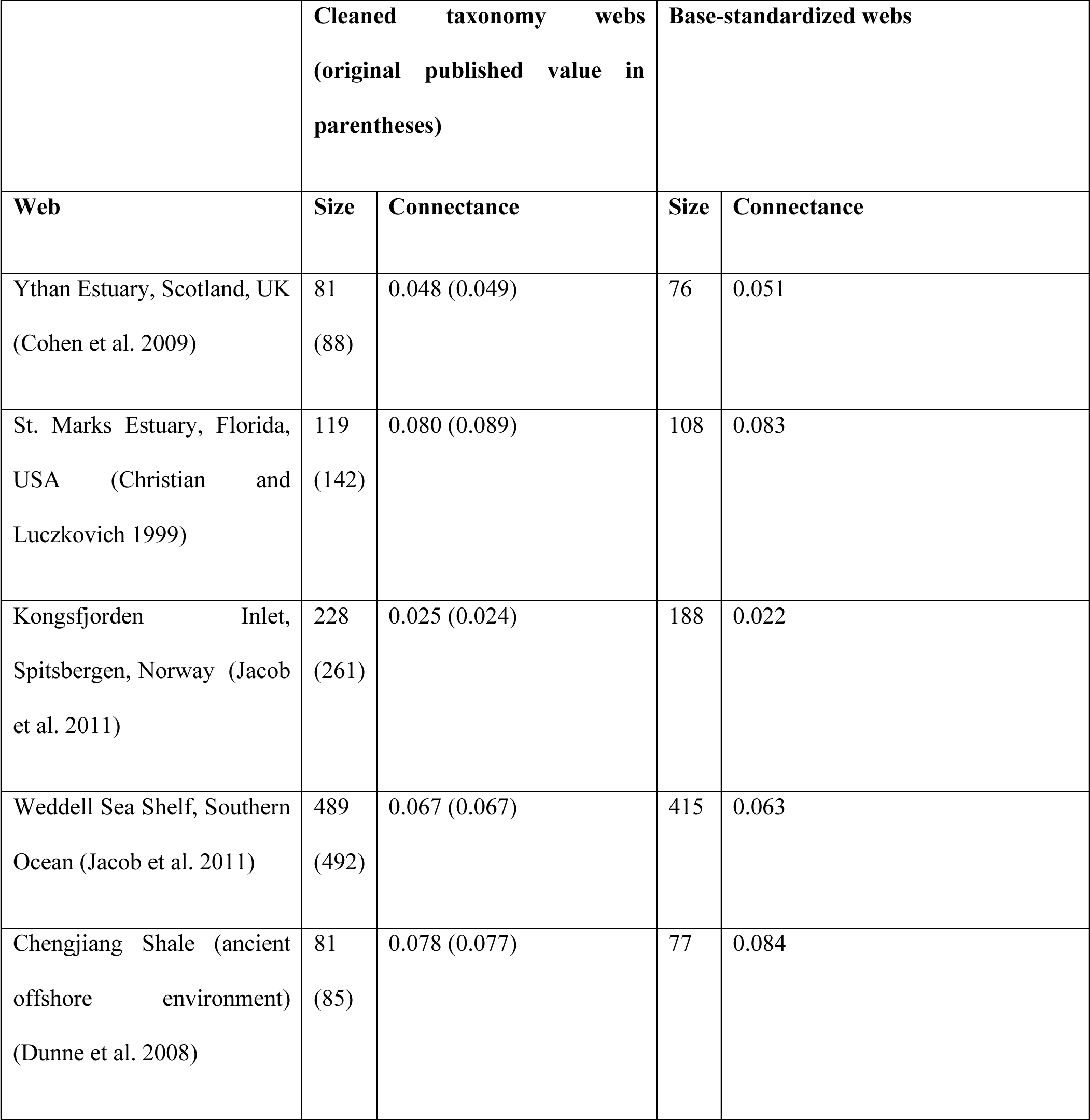
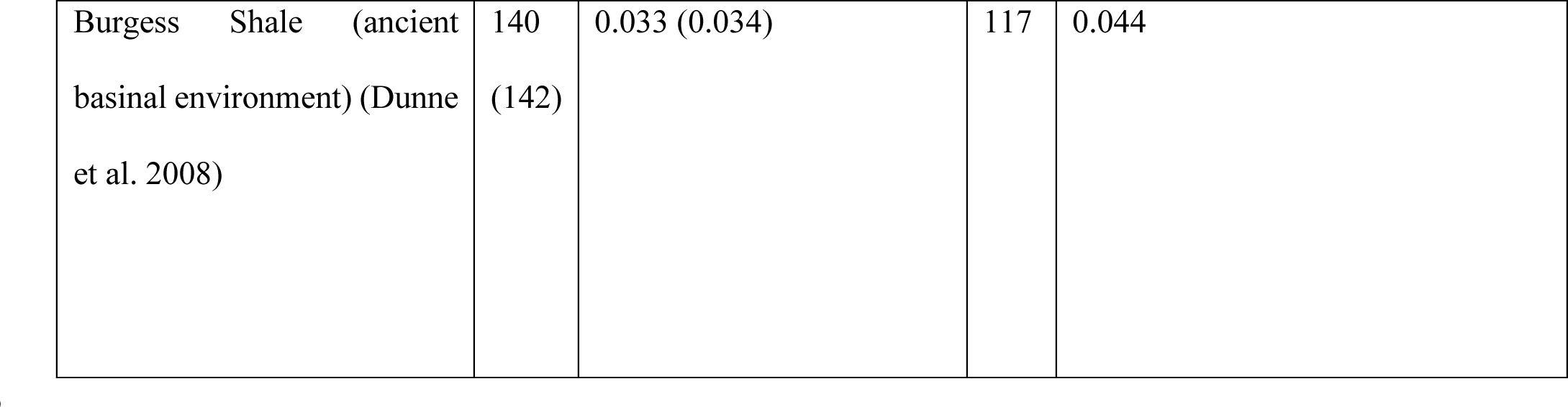
Details of food webs considered in this study.

|  | **Cleaned taxonomy webs**<br>**(original published value in parentheses)** |  | **Base-standardized webs** |  |
| --- | --- | --- | --- | --- |
| **Web** | **Size** | **Connectance** | **Size** | **Connectance** |
| Ythan Estuary, Scotland, UK<br>(Cohen et al. 2009) | 81<br>(88) | 0.048 (0.049) | 76 | 0.051 |
| St. Marks Estuary, Florida,<br>USA (Christian and<br>Luczkovich 1999) | 119<br>(142) | 0.080 (0.089) | 108 | 0.083 |
| Kongsfjorden Inlet,<br>Spitsbergen, Norway (Jacob<br>et al. 2011) | 228<br>(261) | 0.025 (0.024) | 188 | 0.022 |
| Weddell Sea Shelf, Southern<br>Ocean (Jacob et al. 2011) | 489<br>(492) | 0.067 (0.067) | 415 | 0.063 |
| Chengjiang Shale (ancient<br>offshore environment)<br>(Dunne et al. 2008) | 81<br>(85) | 0.078 (0.077) | 77 | 0.084 |
| Burgess Shale (ancient<br>basinal environment) (Dunne<br>et al. 2008) | 140<br>(142) | 0.033 (0.034) | 117 | 0.044 |

#### Taxonomic and trait data

We updated taxon synonyms primarily using the World Register of Marine Species (WoRMS; (Horton et al. 2020)) (see supplemental data). We assigned functional trait attributes to each taxon from three categories after Bambach et al. (Bambach et al. 2007): tiering position relative to the sediment-water interface (six potential states), feeding strategy (four states), and motility (six states). Taxa were assigned the dominant trait if multiple applied (e.g., if a taxon is both a predator and suspension feeder, we selected the primary feeding source based on literature; see supplemental data). Updated taxonomic and life habit assignments were gathered from—and cross-checked between—a series of standard sources (Briggs et al. 1994; Bambach et al. 2007; Xian-Guang et al. 2017; Cirtwill and Eklöf 2018; Horton et al. 2020; Froese and Pauly 2022). We use site-specific body mass values to describe size for modern taxa (Cirtwill and Eklöf 2018) and use length for ancient taxa (given that calculating ancient taxon mass is not feasible; taken from standard sources).

### Primary Producers in ancient food webs

Fossil faunal lists commonly contain no, or very few, non-metazoans and/or primary producers. Fossil surveys commonly focus on one type of fossil such as metazoan body fossils, trace fossils, or plant fossils. For animal fossil assemblages, metazoan taxa are generally identified to a rank between phylum and species, whereas non-metazoans are identified at much coarser resolutions if at all. Thus, to compare ancient food webs and modern food webs, we tested how the removal of non-metazoans from modern food webs impacted apparent food web structure including network and node-level metrics (Sup. Figs. 1-3) and chose a single “base-standardized” approach for all subsequent food-web reconstructions and analyses. In our base-standardized webs, the webs are comprised entirely of metazoans, with all primary producers (and non-metazoans) collapsed into a single basal node.

### Comparing empirical and inferred meta- and realized-food webs

We explored the ability of PFIM to recreate networks in three ways. First, to consider the performance of PFIM with known inputs, we compared model sensitivity across a range of different food web structure scenarios based on three idealized food webs with different proportions of basal, intermediate, and top taxa. Second, we compared the performance of the PFIM against four modern, empirical food webs (i.e., known—rather than inferred—trophic interaction data) where there are associated trait data necessary for reconstructing webs using the model. Specifically, for each assemblage, we compare among empirical webs (i.e., previously published), PFIM-inferred feasible food webs, and PFIM-inferred realized food webs. Third, we compared PFIM-inferred interactions to other sources of interaction data and an alternative interaction inference framework. Networks were compared in terms of (i) network- and node- level metrics (Table 2) and (ii) proportion of correctly identified links.

**Table 2:** Descriptions of network metrics used in this study.

| Metric type | Abbreviation | Metric | Description |
| --- | --- | --- | --- |
| Network structure | C | Connectance | Proportion of possible links in a network that actually occur. Calculated as the number of links (L) divided by number of species (S) squared (i.e. $L/S^2$). |
|  | S | Size | Number of nodes in a graph. |
|  |  | Diameter | Shortest distance between the two most distant nodes of a network, (i.e., the longest path of all shortest paths between each pair of nodes). |
|  | Mean btw. | Mean normalized betweenness centrality | See node position metrics below. |
|  | SD gen. | Standard deviation of normalized in-degree | See node position metrics below. |
|  | SD vul. | Standard deviation of normalized out-degree | See node position metrics below. |
|  | Mean TL | Mean trophic level | See node position metrics below. |
|  | Max TL | Max trophic level | See node position metrics below. |
|  | SOI | Systems omnivory index | Weighted average of consumer OI values (see node position metrics below), where weighting is based on the number of each consumers in-degree (number of prey). |
| | Motif: Lin. ch. | Normalized linear chain motifs | Number of motifs (3-node subgraphs) describing simple linear chains, normalized to $S^2$. |
| | Motif: Omn. | Normalized omnivory motifs | Number of motifs describing predators preying on two taxa at different trophic levels, normalized to $S^2$. |
| | Motif: Ap. comp. | Normalized apparent competition motifs | Number of motifs describing predators preying on two species, normalized to $S^2$. |
| | Motif: Dir. comp. | Normalized direct competition motifs | Number of motifs describing two predators sharing a prey species, normalized to $S^2$. |
| Node position | Btw. | Normalized betweenness centrality | Extent to which a node lies on paths between other nodes. Calculated as the number of shortest paths that pass through a given node and normalized to size of network excluding the node in question ( $S - 1$ ). |
|  | Generality | Normalized in-degree | How many resource nodes a consumer node has. Normalized to the average number of links per node for that network. |
|  | Vulnerability | Normalized out-degree | How many consumer nodes a resource node has. Normalized to the average number of links per node for that network. |
|  | TL | Trophic level | 1 + the weighted average of the trophic levels of its resources. Primary producers are assumed to have trophic level of 1. |
|  | OI | Omnivory index | Variation in trophic levels of the resources of a consumer. |

For the first approach we defined three characteristic faunal lists. In the “standard” web, the faunal list contained two nodes for each functional group (that is, each unique combination of tiering, motility, and feeding traits) and a basal node representing primary production. In the second “top-heavy” web, nodes (representing functional groups) with trophic levels above the median trophic level in the standard web were represented by three distinct nodes while those with trophic levels below the median were represented by a single node including the basal primary production node. In the third “bottom-heavy” web, functional groups with trophic levels below the median in the standard web were represented by three distinct nodes (representing distinct taxa with identical traits), except for a single basal primary production node, while those with trophic levels above the median were represented by a single node. Each resultant feasible food web had the same number of nodes (S = 253) but had a different distribution of trophic levels. We generated corresponding realized webs (n = 1000) for each feasible food web type (i.e., standard, top-heavy, and bottom heavy) and measured network- and node-level metrics to consider how applied link distributions differentially affect food webs with different node distributions.

For the second approach, we assessed the predictive accuracy of PFIM in generating feasible and realized food webs by comparing the PFIM-predicted links to the empirical data. Specifically, we calculated: true positives as the number of links predicted in the inferred feasible (or realized) food web and observed in the empirical web (denoted by “*a”*); false positives as the number of links predicted but not observed (*b*); false negatives as the number of links not predicted but observed (*c*); and true negatives as the number of links neither predicted nor observed (*d*). We summarized these values using three metrics: sensitivity, the proportion of observed presences correctly predicted (a/(a+c)); specificity, the proportion of observed absences correctly predicted (d/(b+d)); and the True Skill Statistic (TSS), a single value of overall model predictive accuracy equally weighting sensitivity and specificity (TSS=[(*ad*−*bc*) ⁄ ((*a*+*c*)(*b*+*d*))], which is also equal to sensitivity + specificity – 1) (Allouche et al. 2006). We also calculate total correct predictions as the percentage of true positives (i.e., a) plus true negatives (i.e., d).

For the final approach, we further delve into an issue left unresolved by the second approach— that is, whether false positives are the result of a true absence of interaction or the absence of observation of an interaction that did occur. To do so, we further analyzed these links by comparing links in the empirical and inferred webs to (i) an online database of trophic interactions (GloBI”) and (ii) a food web inferred utilizing the Allometric Diet Breadth Model (“ADBM”). We focused on the Ythan Estuary web given that it has been re-created using the ADBM (Petchey et al. 2008) and its relatively small size permitted the computationally intensive comparison to GloBI-recorded interactions. We calculated the proportions of true positives, false positives, false negatives, and true negatives as compared to (i) alternative assemblages (i.e., GloBI) and to (ii) the alternative inference method applied to the same taxon list (i.e., ADBM).

Network- and node-level metrics are sensitive to the number of nodes (i.e., number of species) and links (e.g., number of interactions amongst nodes), such that comparisons between webs required normalization. To compare empirical webs with the inferred feasible and realized webs, we normalized webs using null-model food webs generated with the Niche Model (Williams and Martinez 2000). Normalized network-level metrics are presented in terms of effect size, which is calculated as the difference between the target metric value and the null model median metric value (based on 1000 replicates), scaled to a unit variance by dividing by the difference between the upper (or lower) bound of the 95% confidence bound value and the median (depending on whether the value was positive or negative) (e.g., (Dunne et al. 2008)). Once normalized, we refer to webs as “niche-normalized webs”. Effect sizes greater than 1 or less than -1 indicate significant differences between the focal web and the null model.

We attempted to use machine learning models to quantify the predictive power of functional trait data to generate accurate food webs that could be applied to novel data, but we lacked the statistical power necessary to do so (see supplement for more details; Sup. Figs. X,X).

## RESULTS

The three validation methods reveal different aspects of how PFIM performs, and of the relationship between feasible and realized food webs. We discuss the results of each validation method in turn.

### PFIM construction of idealized meta- and realized-food webs

Network metrics successfully differentiate among the idealize PFIM webs with different shapes (e.g., bottom- and top-heavy trophic structures). Resultant metrics differ with the underlying node distribution regardless of whether feasible or realized food webs are studied (Fig. 3). Some metrics that are strongly differentiated in the feasible food webs, like connectance (median ranges from 0.06 to 0.16 in the feasible networks), show minimum differences in the realized webs (median connectance ranges from 0.01 to 0.03), after the imposition of an assumed link distribution. Furthermore, while the distribution of generality (i.e., number of resource items by consumer) values appear similar between web shapes—with lower SD generality displayed by bottom-heavy structures for both feasible and realized food webs—the distribution of vulnerability shows large differences between feasible and realized food webs. Differences between feasible and realized webs are generally of similar magnitude between different web shapes, as indicated by effect size values (in which feasible food web structural metric values were normalized relative to the median and distributions of realized web metric values; Sup. Fig. 4), although bottom heavy webs generally have lower errors than standard and top-heavy webs. Similarly, node-level metrics vary more widely in top-heavy structures, and standard structures, as compared to bottom-heavy structures, both in terms of distributions of node metric values between taxa within food web shapes (i.e., the ranges and averages within webs, Fig. 4 left), but also in terms of the positions of identical taxa between food web shapes (i.e., standard deviation of values across the webs, Fig. 4 right).

**Figure 3:**
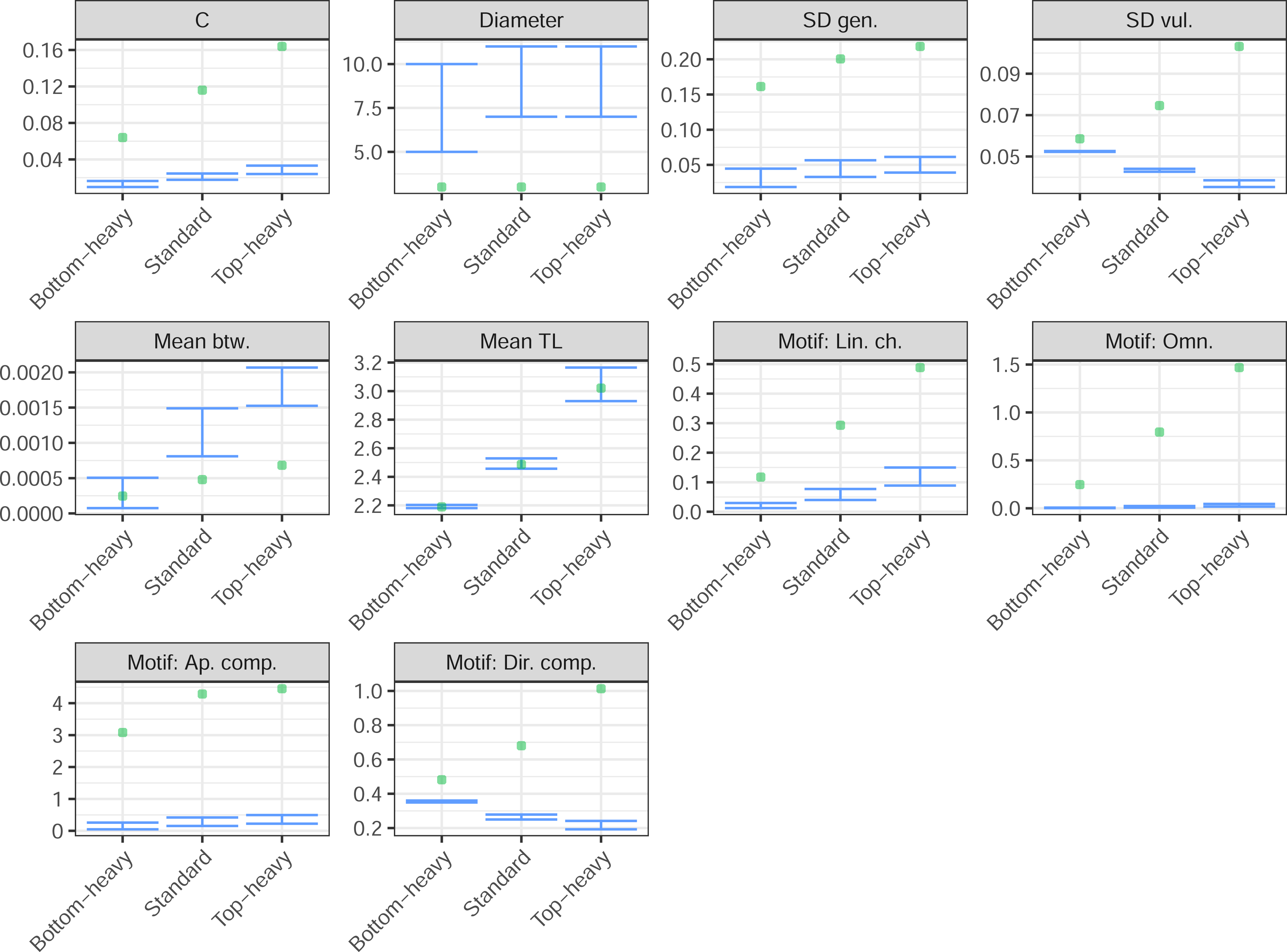
PFIM reconstruction model food webs as assessed using network metrics on the feasible food webs. Network metrics for feasible food webs (green points)—and corresponding realized webs (blue whiskers indicating minimum and maximum values; n=1000)—based on different food web “shapes” including “bottom-heavy” with more basal nodes, “standard” with nodes equally dispersed across all trophic levels, and “top-heavy” with more predatory nodes (see methods for details).

**Figure 4:**
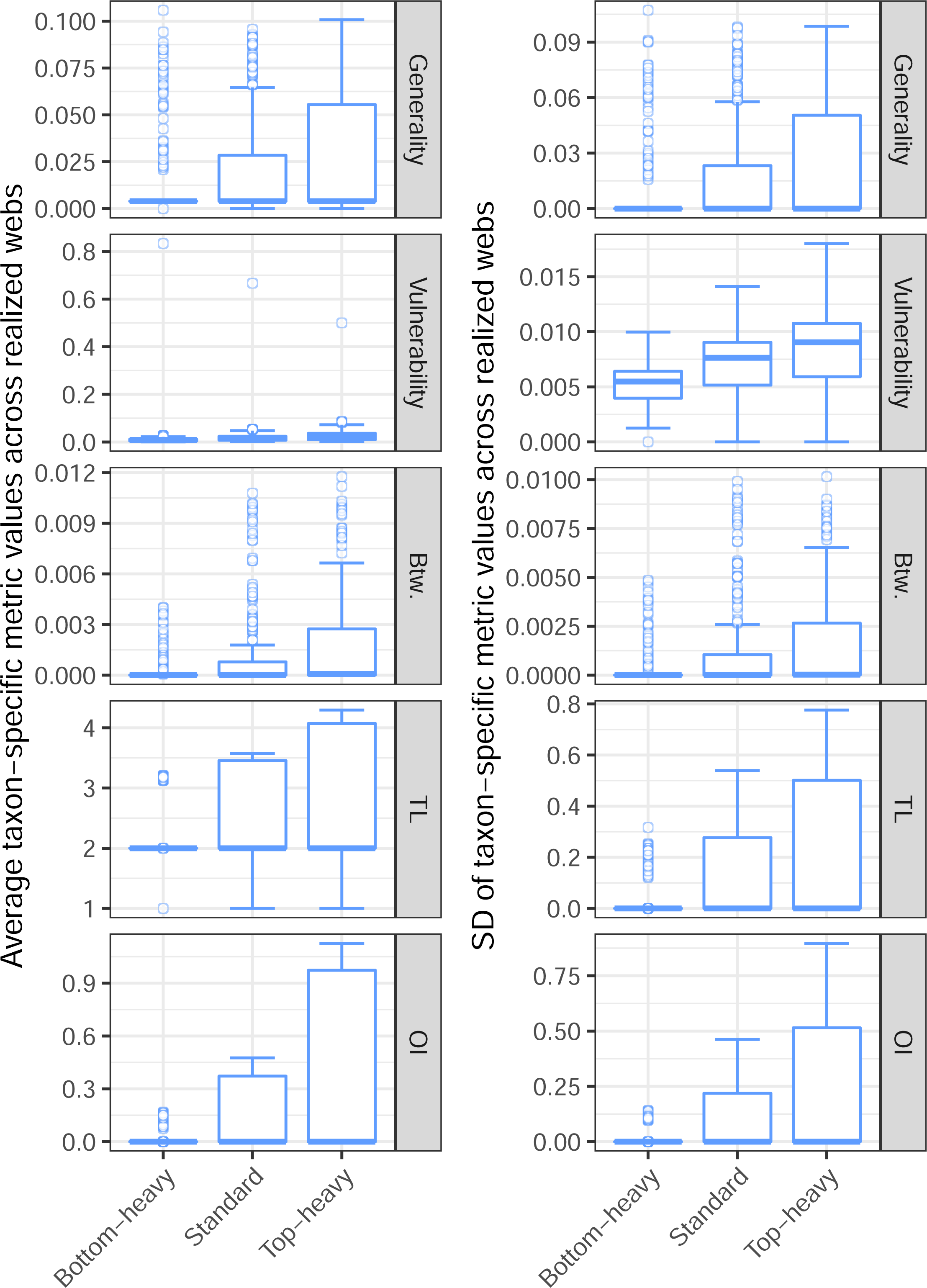
PFIM-generated realized food webs as assessed using node-level metrics (average in left column; standard deviation in right column) of hypothetical shapes (n=1000).

### PFIM versus empirical food webs

Empirical webs and the PFIM-inferred feasible food webs differ in terms of assigned interactions (Fig. 5), network-level structure (Fig. 6), and node-position distributions (Sup. Fig. 5), yet PFIM correctly infers the presence (or absence) of most links within the empirical data. Network- and node-level metrics related to connectance (e.g., mean degree) and the variation in connectance (standard deviation of generality, standard deviation of vulnerability), were distinctly high in the feasible food webs, as were network-level metrics related to chain-length and number, particularly diameter and normalized linear chain motifs (Fig. 6). Differences between empirical and inferred feasible networks generally correlated with overall web size (i.e., the greatest differences in the biggest webs). Normalizing for the connectance-dependence of metrics via the Niche Model corrected some, but not all, of the differences between meta- and empirical webs (Sup. Fig. 6).

**Figure 5:**
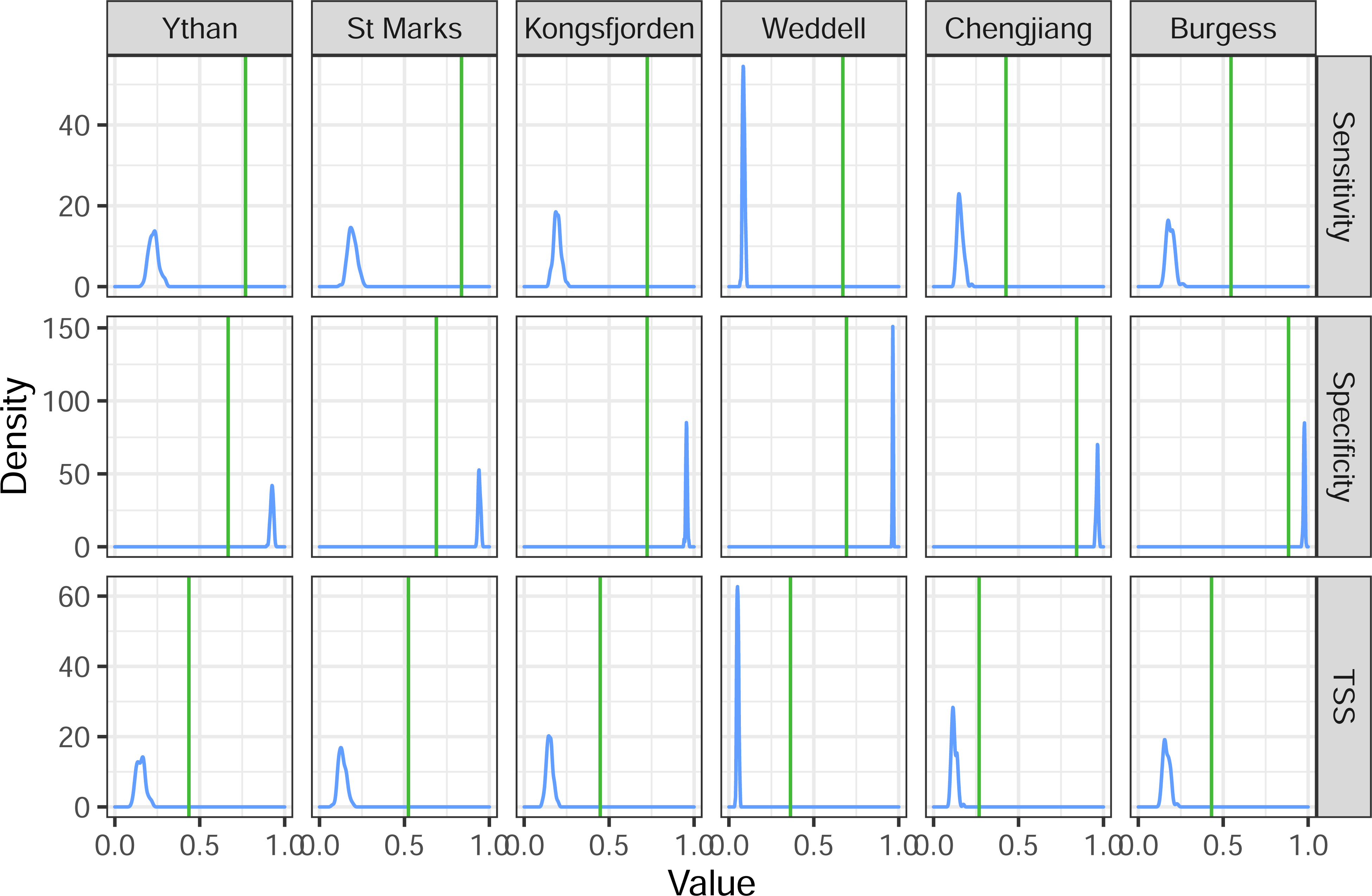
Link prediction accuracy of PFIM feasible (green lines) and realized webs (blue density distributions, 1000 replicates), as compared to the corresponding empirical webs for the four study sites. Sensitivity is the proportion of observed presences correctly predicted (i.e., (true positives)/(true positives + false negatives)). Specificity is the proportion of observed absences correctly predicted (i.e., (true negatives)/(false positives + true negatives)). TSS (True Skill Statistic) summarizes sensitivity and specificity, equally weighting the two (i.e., TSS = sensitivity + specificity – 1). Webs ordered by size (see Table 1).

**Figure 6:**
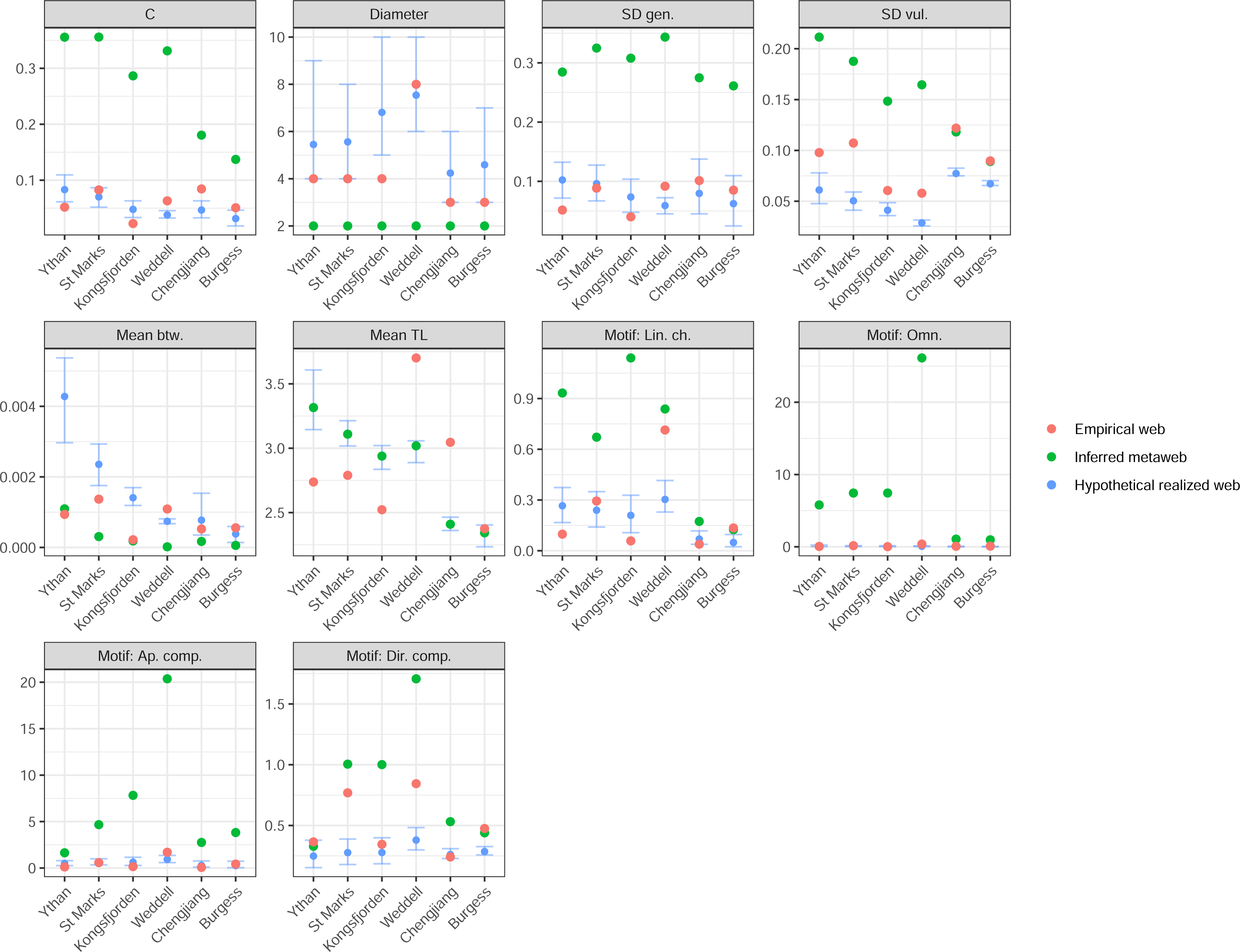
Network-level statistics of empirical, PFIM inferred feasible webs, and PFIM realized webs for the four study sites. Red and green points describe single metric values for empirical and inferred webs, respectively. Blue points describe the mean of metric values of 1000 realized food webs, and the accompanying error bars describe 95% confidence intervals of these values.

In contrast to the feasible food webs, the realized food webs were structurally more similar to empirical food webs (Figs. 6). At least two of the four empirical webs fell within the range of values for realized webs, including diameter, maximum trophic level, and three motif metrics (omnivory, apparent competition, and direct competition) (Fig. 6, Sup. Fig. 6). Although other metrics were largely non-overlapping between the empirical and hypothesized webs, the difference in the case of several metrics (connectance, mean degree, standard deviation of generality, and SOI) was on the order of variation observed among empirical webs from different localities. Empirical and realized webs differed most in mean trophic level and the standard deviation of vulnerability (Fig. 6).

PFIM-inferred feasible food webs correctly infer ∼70% of interactions—or lack thereof—found in empirical food webs (i.e., true positives plus true negatives; Table 3). For realized interaction webs, PFIM correctly infers ∼90% of interactions (i.e., true positives plus true negatives; Table 3). In the feasible food web, correctly predicted absences (true negatives) of consumer-resource trophic interactions accounted for 63-71% of possible links, correctly predicted observations (true positives) of interactions accounted for 2-7% of possible links, and inferred but not recorded observations (false positives) accounted for 27-32% of possible links (Table 3). Less than 2% of possible links were recorded in the empirical web as occurring but were not inferred in the feasible food web (false negatives). For the hypothesized realized web, the link predictions (averaged across 1000 replicates) across the four webs ranged 86-90% for true negatives, 1-2% for true positives, 3-7% for false positives, and for 2-7% false negatives (Table 3). Inferred feasible food webs performed much better than realized webs in terms of correctly predicting present interactions (i.e., sensitivity; Fig. 5), whereas realized webs performed marginally better than feasible food webs in terms of correctly predicting absent interactions (i.e., specificity; Fig. 5). As a result, TSS values of inferred feasible networks were higher, falling between 0.36 and 0.52, compared to realized webs (0.05-0.15; Table 3).When size criteria were removed (i.e., defining feasible interactions based on motility, feeding, and tiering only), TSS values of inferred feasible webs decreased (Ythan = 0.27, St Marks = 0.42, Kongsfjorden = 0.42, Weddell = 0.36), largely due to an increase in the proportion of false positives compared to true negatives.

**Table 3:** Breakdown of link composition and true skill statistic (TSS) scores of inferred feasible and realized webs, both as compared to the corresponding empirical web. Of the 1873 false positives (representing 32% of possible feasible interactions) recorded in the Ythan Estuary web, 1480 (79% of 1873) were recorded only by PFIM, 118 by PFIM and the ABDM (6%), 256 by PFIM and GloBI (14%), and 19 (1%) were recorded by all three datasets (i.e., PFIM, ABDM, and GloBI).

| **Web** | **Total number**<br>**of possible links** | **Type** | **True**<br>**positives**<br>**(a)** | **False**<br>**positives**<br>**(b)** | **False**<br>**negatives**<br>**(c)** | **True**<br>**negatives**<br>**(d)** | **TSS** |
| --- | --- | --- | --- | --- | --- | --- | --- |
| Ythan | 5776 | Feasible | 4% | 32% | 1% | 63% | 0.44 |
|  |  | Realized | 1% | 7% | 4% | 88% | 0.15 |
| St Marks | 11664 | Feasible | 7% | 29% | 1% | 63% | 0.52 |
|  |  | Realized | 2% | 5% | 7% | 86% | 0.13 |
| Kongsfjorden | 35344 | Feasible | 2% | 26% | 1% | 71% | 0.45 |
|  |  | Realized | 1% | 4% | 2% | 93% | 0.15 |
| Weddell | 172225 | Feasible | 4% | 29% | 2% | 65% | 0.36 |
|  |  | Realized | 1% | 3% | 6% | 90% | 0.05 |

### Comparisons with alternative sources of interaction data

In our analyses of false positives in the Ythan Estuary web, we found that 21% of these interactions were recorded by alternative datasets (Table 3), of which 14% were observed in other assemblages (i.e., GloBI) and another 7% were inferred to exist using the ADBM, with 1% indicated by both GloBI and the ADBM. When false positives recorded by GloBI and the ADBM (i.e., indicating that these interactions are feasible and realizable; 393 interactions; see Table 3 and caption) are instead considered as true positives, the TSS of the PFIM-inferred Ythan feasible food web rises to 0.62 (from 0.44; Table 3).

### Reconstructing Cambrian food webs using PFIM

When analyzing the prediction accuracy ancient webs based on functional trait information, we find that sensitivity is lower for both the Burgess and Chengjiang assemblages. Although the TSS value for Chengjiang is lower than values for modern assemblages, the Burgess falls within the range of values for those assemblages. When considering realized web structures, sensitivity, specificity, and TSS values are similarly distributed between ancient and modern assemblages.

Comparisons of the network-level statistics between ancient and modern assemblages (both in terms of metawebs and realized webs) show that the Burgess and Chengjiang networks have relatively low connectance, SD vulnerability, and mean TL values. While node-level statistics largely mirror this trend, taxa in the ancient webs display higher generality and lower vulnerability when compared to taxa in modern webs.

## DISCUSSION

The Paleo Food web Inference Model (PFIM) provides a simple platform in R to build trophic networks—both feasible and realized food webs—based on the data available in the fossil record. This approach provides an platform for studies of ancient food webs building on previous work establishing the relationship between functional traits and trophic interactions (e.g., (Eklöf et al. 2013; Cirtwill and Eklöf 2018); also see supplemental analysis of functional structure of networks studied herein; Sup. Figs. 7,8) and the utility of inference modeling in modern and ancient ecosystems (e.g., (Beckerman et al. 2006; Roopnarine 2006; Petchey et al. 2008)).

Although PFIM networks are imperfect replications of published food webs—which, themselves, are imperfect recorders of trophic interactions and food web structure (Polis 1991)—the three validation procedures carried out here highlight the potential and limitations of this method.

Using functional trait data and a set of consumer-resource interaction criteria alone we correctly predicted ∼70% of interactions—or lack thereof—found in four empirical food webs (i.e., true presence and true absence of interactions; Table 3). This figure rose to ∼90% when considering realized interaction food webs for those same ecosystems. However, for the PFIM feasible food webs, ∼29% of all interactions predicted by the inference model were not found in the empirical web (false positives) –these interactions may represent unsampled, unrealized, or unfeasible links. Indeed, our in-depth analyses of the Ythan Estuary web, indicate that some of the interactions inferred but not observed may be unsampled, given that 21% of false positives were recorded in other datasets (i.e., GloBI and ADBM) (Table 3). Intriguing though this finding is, this relatively high rate of under-sampling may be exclusive to the Ythan Estuary web, which appears less well resolved than other assemblages considered here (e.g., a quarter of nodes are birds, possibly indicating an underrepresentation of aquatic taxa). Despite these shortcomings, the Ythan web remains one of the few well-studied systems with data suitable for comparisons with PFIM-inferred networks (as discussed in methods).

Only a small fraction of links (∼1%) found in the empirical food webs were not inferred in the feasible food webs, suggesting that the many “false positives” are predominantly unrealized or unsampled in the empirical data, rather than truly unfeasible interactions. Unfeasible links could include cases allowed by our simplified criteria for interaction that are functionally impossible. For instance, a predator that cannot break shells could not eat bivalves still might well meet all four interaction criteria in PFIM. Generally, however, the relatively high predictive accuracy of the PFIM feasible food webs indicates the utility of the simplified set of interaction criteria in cases where little interaction information is available, as in fossil assemblages (see supplement for additional considerations).

Across deep time, the realization of links between taxa is likely to vary substantially due to changing realized niches, evolution, and migration (Veloz et al. 2012), which can vary over just a few generations (Kraan et al. 2013). Thus, the over-connectance of PFIM feasible food webs relative to the empirical food webs may reflect the full potential for re-wiring on evolutionary timescales. Even after niche-normalization (Sup. Fig. 6), the PFIM feasible food webs were structurally distinct from the empirical and hypothesized realized food webs. In other words, the feasible food webs differ from the empirical food webs for reasons beyond the number of links, and these structural differences may reflect the eco-evolutionary potential for changes in intraspecific interactions. Much remains to be learned about how and why a feasible interaction might be realized in modern assemblages (Poisot et al. 2015), let alone ancient ones. On evolutionary time scales, consumers can adjust their diets and foraging behavior in response to external changes (Gilljam et al. 2015) and this flexibility is thought to confer stability upon systems (Henri and Van Veen 2016). PFIM feasible food webs can be used to directly consider how interactions—and trophic structure—may change across evolutionary timescales (e.g., food web re-wiring) in response to events like evolutionary innovations and abrupt environmental perturbations.

The idealized web-shapes highlight potential eco-evolutionary differences among food webs for rewiring due to changes in the distribution of nodes through time. Our three idealized food webs differed in the distribution of nodes among trophic level from an even distribution (“standard”) to most nodes occurring at in low trophic levels (“bottom-heavy”) or high trophic levels (“top-heavy”). This difference in node distribution led to distinct differences in feasible and realized food webs with realized networks for progressively top-heavy web-shape showing the greatest variation in wiring at the node- and network-level (Figs. 3,4; Sup. Fig. 4). This exploration of idealized web-shapes highlights two important aspects of the PFIM approach for considering food webs through time. First, the relationship between feasible and realized food webs is strongly dependent on the underlying distribution of nodes, as well as the assumed link distributions. Second, long-term trends in the occupation of functional modes of life through time (Bush and Bambach 2011), with trends towards increased energetics and body sizes (Finnegan et al. 2011), should be accompanied by commensurate changes in the potential for food web rewiring.

PFIM provides a flexible framework for generating a series of realized food webs using different characteristic distributions of links among species (e.g., power–law, exponential, and uniform degree distributions (Dunne et al. 2002*a*)). By default, PFIM generates hypothetical realized food webs with exponential-power law in-degree distributions based on previous research on fossil food webs (Roopnarine et al. 2007), a link distribution type similar to those found in modern and ancient communities (Dunne et al. 2002*a*, 2002*b*, 2008). This approach generated a series of food webs with node- and network level characteristics like the empirical food webs (Fig. 6), predominantly due to the imposed exponential power-law distribution of interactions.

Although hypothesized realized food webs may match well with modern empirical food webs (e.g., (Roopnarine and Dineen 2018)), the assumption of constant link-distributions through time—implied by applying the same distribution to modern and ancient assemblages—may not hold. Multiple different link distribution hypotheses can thus be used to address questions related to changes in the structure, energy flow, and stability of food webs through time given the full range of link distributions that may have characterized the ancient food webs.

PFIM, based on a simple set of traits, performs comparably to alternative inference methods requiring more data on the taxa for which interactions are being predicted. For instance, a Niche

Model style approach, parameterized based on known predator-prey size ratio data, generates networks with TSS values ranging from 0.13 to 0.76 (as compared to the 0.36-0.52 values for PFIM feasible webs), when applied to Mediterranean pelagic fishes (Gravel et al. 2013).

## CONCLUSIONS

Developing PFIM highlighted several best practices for constructing and comparing ancient food webs, given data commonly available to paleoecologists, and potential avenues for future model development. First, when considering comparisons between ancient food webs, we recommend the use of base-standardized metazoan networks, given that non-metazoans and metazoans are rarely sampled to the same degree in the same deposit (see supplement for details). Second, feasible and realized webs are suited to exploring different issues related to the structure and dynamics of food webs through time. Feasible food webs allow for eco-evolutionary considerations related to trophic rewiring as well as a link-distribution-agnostic approach to the history of trophic interactions. A feasible interaction perspective may be particularly fitting for fossil assemblages given that time averaging combines individuals that may never have co-occurred (Kidwell and Holland 2002) thereby preserving communities already representative of eco-evolutionary timescales. Third, not all fossil deposits are created equal. PFIM is expected to perform best in reconstructing ancient food webs when using the relatively comprehensive fossil data available from Konservat-Lagerstätten (Shaw et al. 2021*a*, 2021*b*). When other deposit types are used the effect of biases on assemblage composition, such as the lack of soft-bodied taxa, networks will appear less stable, with less predation and an overrepresentation of generalist consumers (Shaw et al. 2021*b*). On top of preservation biases, human biases, such as the preferential sampling of enigmatic clades and historical biases in sampling locations (e.g., (Raja et al. 2021)), also skew interpretations of ancient biodiversity.

Here we show how a simple set of organismal traits—motility, feeding, tiering, and size—can be used in PFIM to generate feasible and realized food webs for modern and ancient trophic systems. PFIM is a pragmatic attempt to balance confidence, given the constraints on data from the geological record, and utility, and presents the opportunity to consider ecosystem dynamics more robustly across deep time in data-limited assemblages.

## ACKNOWLEDGEMENTS

We thank S. Allesina, D.E.G. Briggs and A. Eklöf for insightful conversations. We also thank O.L. Petchey for supplying the ADBM food web for Ythan Estuary. J.O.S. was supported by funding from the Yale Institute for Biospheric Studies, the Yale Franke Program in Science and the Humanities, and the Santa Fe Institute.

## AUTHOR CONTRIBUTIONS

J.O.S. developed methods, collected data, analyzed data, developed software, interpreted results, wrote the original draft, and edited subsequent drafts of this paper. A.M.D. developed methods, interpreted results, and reviewed and edited drafts. A.P.B. developed methods, interpreted results, and reviewed and edited drafts. J.A.D. interpreted results and reviewed and edited drafts. P.M.H. developed methods, interpreted results, reviewed and edited drafts, and advised the lead author.

## DATA AVAILABILITY

The final versions of all data, code, and supplementary information in this paper will be uploaded to Dryad and assigned a DOI. Updates to code will be posted to a permanent GitHub link. During review data will be made available to reviewers.

