## Supplement for "A framework for reconstructing ancient food webs using functional trait data"

### Data collection and editing

#### Including non-metazoans in ancient metazoan-focused food webs

To understand how best to compare ancient food webs to each other and to modern food webs we tested how the removal of non-metazoans from metazoan-focused food webs impacted apparent food web structure. These tests were carried out because metazoan and non-metazoans fossils are typically collected in different deposits, making it difficult to reconstruct a full-food web for any given locality and time.

To compare the structures of webs with and without non-metazoans we had to account for systematic differences in network-level metrics dependent on the number of nodes (representing organisms) and links (representing interactions between two nodes) in a particular web (with the metazoan-only webs having fewer of both). To do this we normalized webs by comparing them with simulated null model food webs generated with the "niche model" (i.e., generating "niche-normalized webs"; see main text for more in-depth description of niche normalization).

We compared network- (Sup. Figs. 1,2) and node-level (Sup. Figs. 3) metrics between five versions of the same food web edited based upon different qualities and resolutions of data:

1. Original published web, with node names updated to reflect updated or accepted taxonomic designations.
2. Cleaned published web, a version of the original web where metazoan nodes have been removed if they lacked complete functional data (i.e., tiering, feeding, motility, and size classifications) or family-level or below taxonomic data.
3. Multiple-basal-nodes web, a version of the "cleaned published web" in which non-metazoan nodes were grouped by high-ranked Linnean groups (mostly at the kingdom-level).
4. Single-basal-node web, a version of the "cleaned published web" web in which all non-metazoans were grouped into a single "basal node".
5. Metazoan-only web, a version of the "cleaned published web" web in which all non-metazoans were removed.

The different webs had slightly different sizes (and thus connectance), due to the removal of non-metazoan nodes. Given the systematic covariation of network metrics with size and connectance, we compared standardized webs to respective niche model webs to generate model error values. Model error values were similar between web types (Sup. Fig. 2), except for the metazoan-only webs, indicating that the small differences in size and connectance can account for the small differences in network metrics between different forms of the same web.

Given concerns over the quality of recording of non-metazoan taxa in fossil faunal lists, we chose to utilize single-basal-node web as the basis of comparisons. This obviates the need for the inclusion of individuals (i.e., cleaned published webs) or kingdom-grouped (i.e., Multiple-basal-nodes web) non-metazoan taxa. We recognize the obvious short-comings of this approach including the fact that not all non-metazoans are basal in a food webs, but find it to be a pragmatic solution for now to the undersampling of basal taxa in fossil food webs. For the remaining analyses we focus on comparing food webs inferred using PFIM to single-basal-node style webs, herein referred to simply as the "empirical webs".

### Analyses

#### Quantifying relationship between functional traits and food web structure

To further assess relationships between functional traits and food web structure we analyzed how ecospace occupation (as informed by life habit and size data) varied with network-level metrics and the distributions of node-level metrics in the empirical webs.

First, using node-level metrics we compared how a taxon's functional traits varied with its individual position in the food web. Second, we compared assemblage-specific functional diversity metrics (calculated using (Novack-Gottshall 2020)), which describe the range, evenness, and dispersion of traits within an assemblage, with network-level metrics.

**Functional traits as predictors of trophic interactions**

In the empirical food webs (i.e., base-standardized versions), node-position varied systematically with motility, tiering, feeding, and size. For instance, fast taxa were consistently positioned at higher trophic levels than sessile taxa (Sup. Fig. 7). These systematic variations indicate that the functional trait categories in this study (motility, tiering, feeding, and size) are useful predictors of node-position.

**The functional trait structure of food webs**

Certain combinations of motility, tiering, and feeding traits were represented more than others across the analyzed empirical food webs. For instance, all webs showed an abundance of fast, pelagic, predators compared to a lack of slow, shallow-infaunal, predators (Sup. Fig. 7). Functional diversity metrics (e.g., evenness; Sup. Fig. 8) vary across the webs.

#### Relaxing interaction criteria

When generating a PFIM-inferred feasible food web we used a rule stating that all four interaction criteria (motility, tiering, feeding, and size) must be met between two taxa for an interaction to be deemed feasible. This rule could be relaxed, for instance, by permitting an interaction to be deemed feasible if it meets three of the four interaction criteria regardless of what criteria is unmet. Rather than permitting total flexibility in which interaction criteria are and aren't met, one could relax requirements based on the apparent importance of different variables in predicting (for example, variable importance according to statistical analyses, as shown in our use of machine learning models).

By relaxing interaction criteria requirements, PFIM infers many more links among taxa in an assemblage. This relaxation is likely to generate many functionally impossible links. For instance, if the tiering rule is not met but all three others (e.g., motility, feeding, and size), infaunal and pelagic taxa may appear to easily interact, which is relatively rare in real life. When size criteria were removed (i.e., defining feasible interactions based on motility, feeding, and tiering only), TSS values of inferred feasible food webs decreased (Ythan = 0.27, St Marks = 0.42, Kongsfjorden = 0.42, Weddell = 0.36; as compared to Table 3), largely due to an increase in the proportion of false positives compared to true negatives.

Criteria are broadly applicable to marine metazoans, such that they are relaxed by the generalist nature of trait designations (e.g., "surficial feeders" representing multiple life habits including grazing and mining). The relatively high predictive accuracy of the PFIM feasible food web indicates the utility of the simplified set of interaction criteria in cases where little interaction information is available, as in fossil assemblages. For instance, defining a consistent predator-prey size ratio to be applied throughout time and space is difficult. The size ratios of predator to prey changed substantially over the past ~500 million years, as evidenced by an order of magnitude increase in the relative size of predatory drilling holes in fossil marine shells (Klompmaker et al. 2017). Furthermore, although predator-prey size ratios can be consistent at a global scale (Brose et al. 2019), many individual interactions do not conform to expectations, as indicated by the food webs used herein (e.g., Sup. Fig. 9).

#### Comparing link prediction across PFIM, ADBM, and GloBI

The Allometric Diet Breadth Model food web for the Ythan Estuary was generated by Owen Petchey (personal communications) using methods described in a previous paper (Petchey et al. 2008) applied to interaction data from Cirtwill and Eklöf (Cirtwill and Eklöf 2018). The ADBM-generated food web is included as part of the supplementary data (see supplemental data).

These data were compared to interactions recorded in GloBI. This permitted us to consider how the realization of interactions can vary spatially in the modern. GloBI interaction data was accessed using the rglobi package in *R* and is included as part of the supplementary data (see supplemental data). We generated a list of all possible consumer-resource combinations in the Ythan web and then searched for observed instances of each potential interaction in GloBI.

#### Predicting interactions using machine learning

To quantify the relative power of different functional trait categories in predicting interactions we used an automated machine learning approach ("autoML"). Such machine learning models allow us to simultaneously assess the importance of different variables and generate models that can be applied to novel data to predict interactions in novel assemblages (e.g., (Shaw et al. 2021)). The automated approach allowed us to test several types of machine learning models simultaneously, each with their own strengths, to consider how interactions and dependent variables are structured. Using the "autoML" function of the *R* package h2o (Aiello et al. 2016) we generated two types of machine learning models—Distributed Random Forest (DRF) and Gradient Boosting Machine (GBM)—which were subjected to automatic parameterization and tuning. The most predictive of the models was selected for subsequent analyses. Other machine learning models allow a finer breakdown of variable importance (e.g., the importance of particular levels within a variable). However, this made interpretation of variable importance even more difficult and emphasized the "black box" nature of machine learning models (i.e., it's difficult to tell why a model inferred an interaction or not). Furthermore, given how site-specific our models were, the coarser descriptions of variables (i.e., the importance of a variable overall, rather than specific levels within it) allowed a more robust interpretation of variable importance spanning assemblages, which is more applicable to other assemblages than the results of models producing finer variable breakdowns.

We used the life habits and sizes of consumers and resources (i.e., the data shown in Sup. Fig. 7) to predict the presence or absence of an interaction between all possible pairs of taxa in each assemblage. We created four groups of models: (i) five models separately built and tested using each web using all variables (i.e., consumer life habits, resource life habits, consumer size, and resource size; see web-specific models on Sup. Figs. 10,11); (ii) one model built and tested using all webs and all variables ("All", Sup. Figs. 10,11); (iii) as with (ii) except consumer and resource size variables were excluded ("Excl. size", Sup. Figs. 10,11); (iv) as with (ii) except the assemblage identifier was removed, such that the model did not know which interactions belonged to separate assemblages ("Excl. web", Sup. Figs. 10,11). Models were built using "training data", a random sample of 80% of the data, and tested using the remaining 20%, which allowed us to assess predictive accuracy.

A machine learning model utilizing all available data (pairwise interactions for all five assemblages, and accompanying consumer and resource life habits and size; "All", Sup. Figs. 10,11) identified three consumer variables as the most important—feeding mode, followed by motility, followed by size. The importance of these variables, combined with the importance of resource tiering, confirm the significant of basic tenets of foraging theory in accounting for common functional constraints (i.e., feeding and motility), encounter probability (i.e., motility and tiering), and predator size.

When excluding consumer and resource size information ("Excl. size", Sup. Figs. 10,11) remaining variable importance values change little, except for the increased importance of consumer feeding mode, indicating a close link between feeding mode and size in the food webs studied here. Similarly, when the assemblage ID variable is removed ("Excl. web", Sup. Figs. 10,11), remaining importance values do not change significantly, except for a slightly increased importance of consumer size—this is likely indicative of the different distributions of body sizes between the four food webs.

Site-specific models (i.e., Ythan, St Marks, Kongsfjorden, and Weddell as shown in Sup. Figs. 10,11) show variable unique features. For instance, resource taxon size is the most important variable for predicting interactions in the Ythan food web, yet it is generally ranked low for the other three food webs. Similarly, consumer taxon size is ranked highly for both the Ythan and Kongsfjorden webs, but is ranked lower for the St Marks and Weddell food webs. The unique trends in variable importance for each individual web may indicate that different functional and ecological constraints are structuring the different systems. When site-specific models are applied to the corresponding food web (i.e., the Weddell model built on 80% training data used to predict the 20% test data), predictive accuracy is relatively high (as indicated by TSS values, Sup. Fig. 11). Given the relatively small dataset considered here, further work is needed to evaluate whether functional constraints span systems, or whether they are site-specific as indicated here.

The uniqueness of trends, and the relatively small amount of data considered herein, means that we were unable to produce a model that can be applied more broadly. For instance, a model built on any three of four webs does not have sufficient predictive accuracy when applied to the remaining web. This is confirmed by the low predictive accuracy (e.g., TSS values) of models spanning multiple webs (see the "All", "Excl. size", and "Excl. web" models on Sup. Figs. 10,11).

With more input data (i.e., more interaction data paired with functional trait information and multiple assemblages) machine learning models may become powerful enough to predict interactions in other assemblages based on functional trait data. Unfortunately, the lack of high-resolution food webs (e.g., networks spanning multiple clades and resolved to a low enough level to consider functional traits) currently precludes the generation of such models.

#### Supplementary figure captions

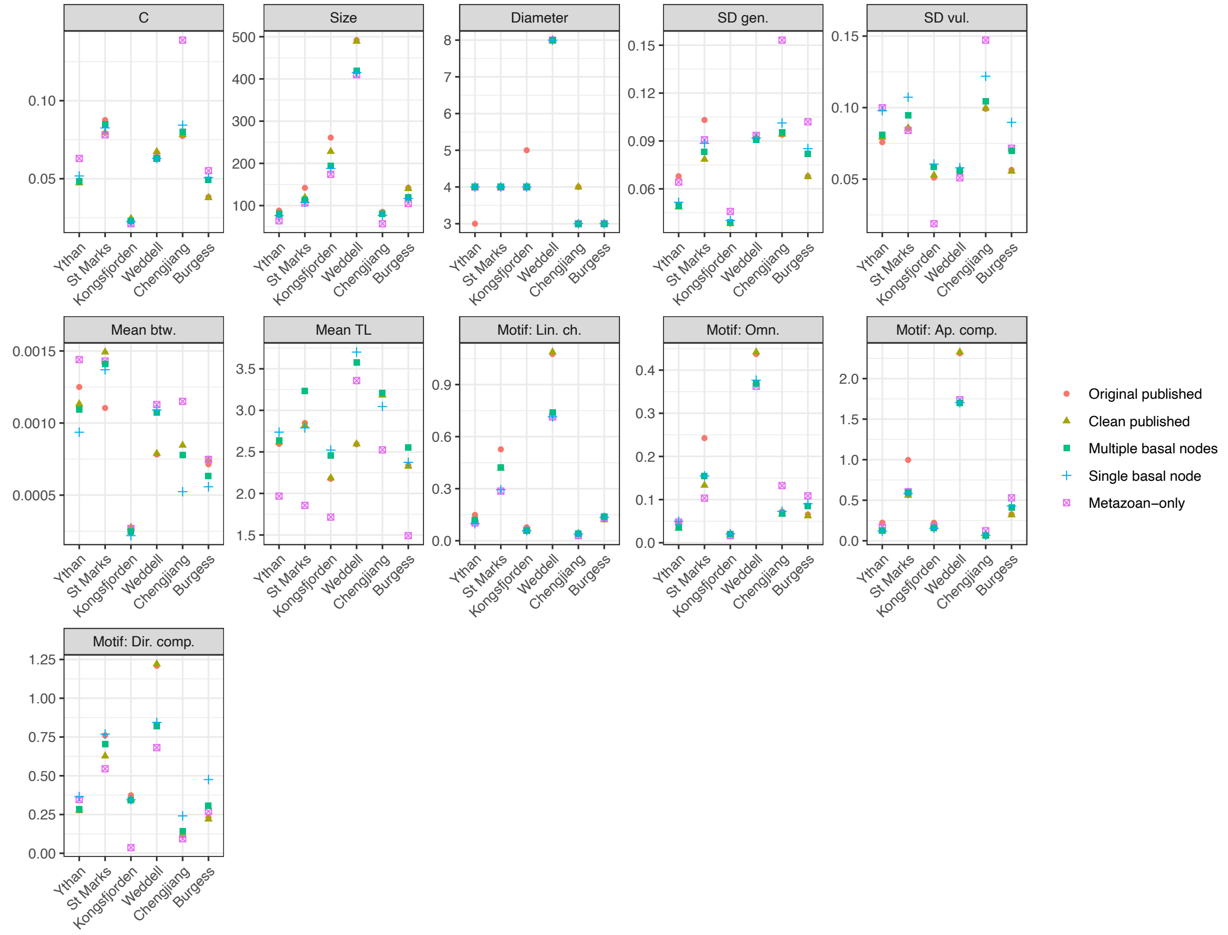

Supplementary Figure 1: Network-level metrics for various edited versions of the empirical food webs considered in this study: original published webs, cleaned published webs, multiple-basal-nodes webs, single-basal-node webs, metazoan-only webs (see supplemental methods for details regarding each version type). Web types ii-v represent webs with similar taxonomic compositions to fossil faunal lists, with non-metazoan taxa coarsely identified or not recorded at all. Webs ordered by size.

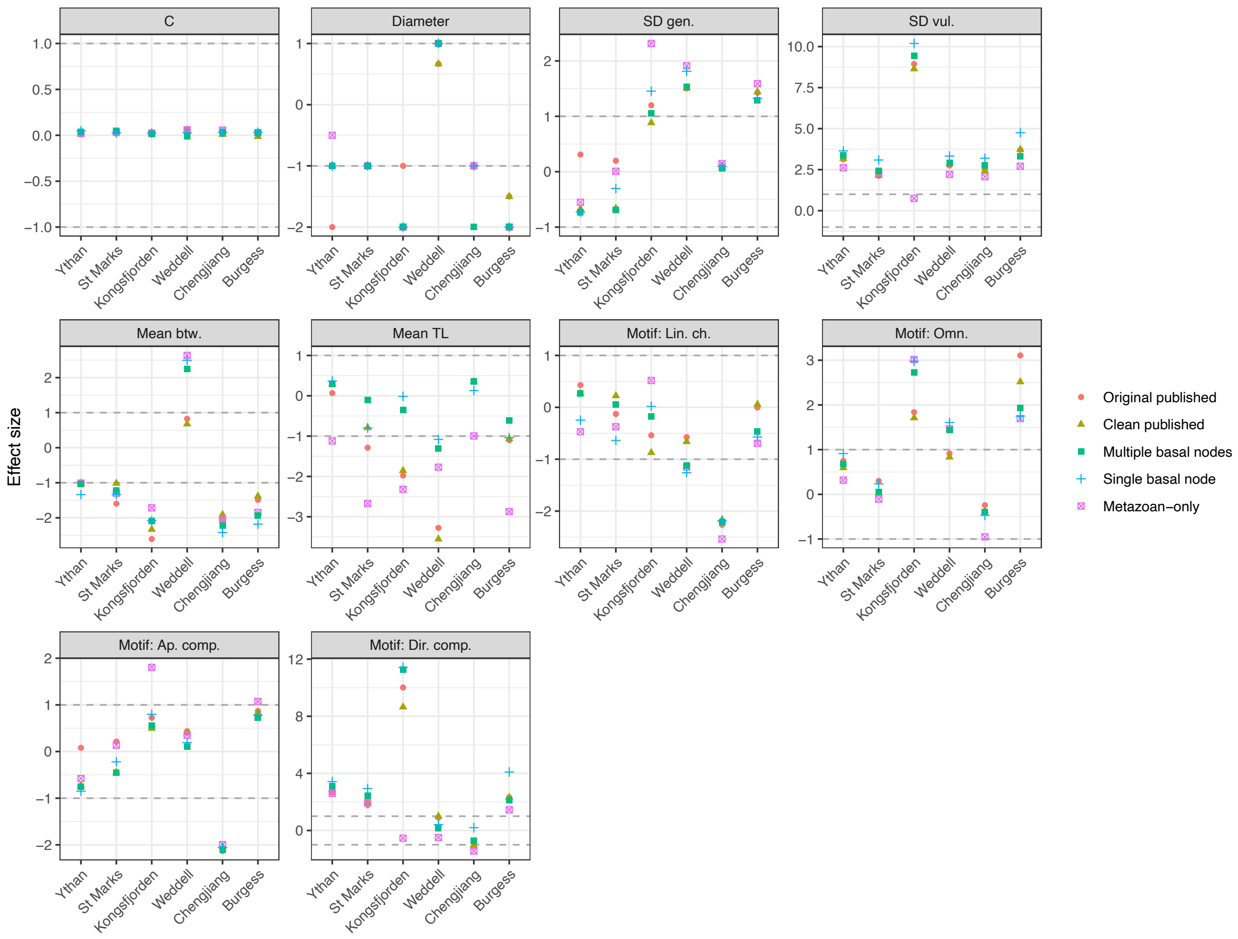

Supplementary Figure 2: Network-level metrics of niche-normalized empirical food webs for various edited versions of the empirical food webs considered in this study: original published webs, cleaned published webs, multiple-basal-nodes webs, single-basal-node webs, metazoan-only webs (see supplemental methods for details regarding each version type). Model error of -1 and +1 indicated by grey horizontal lines.

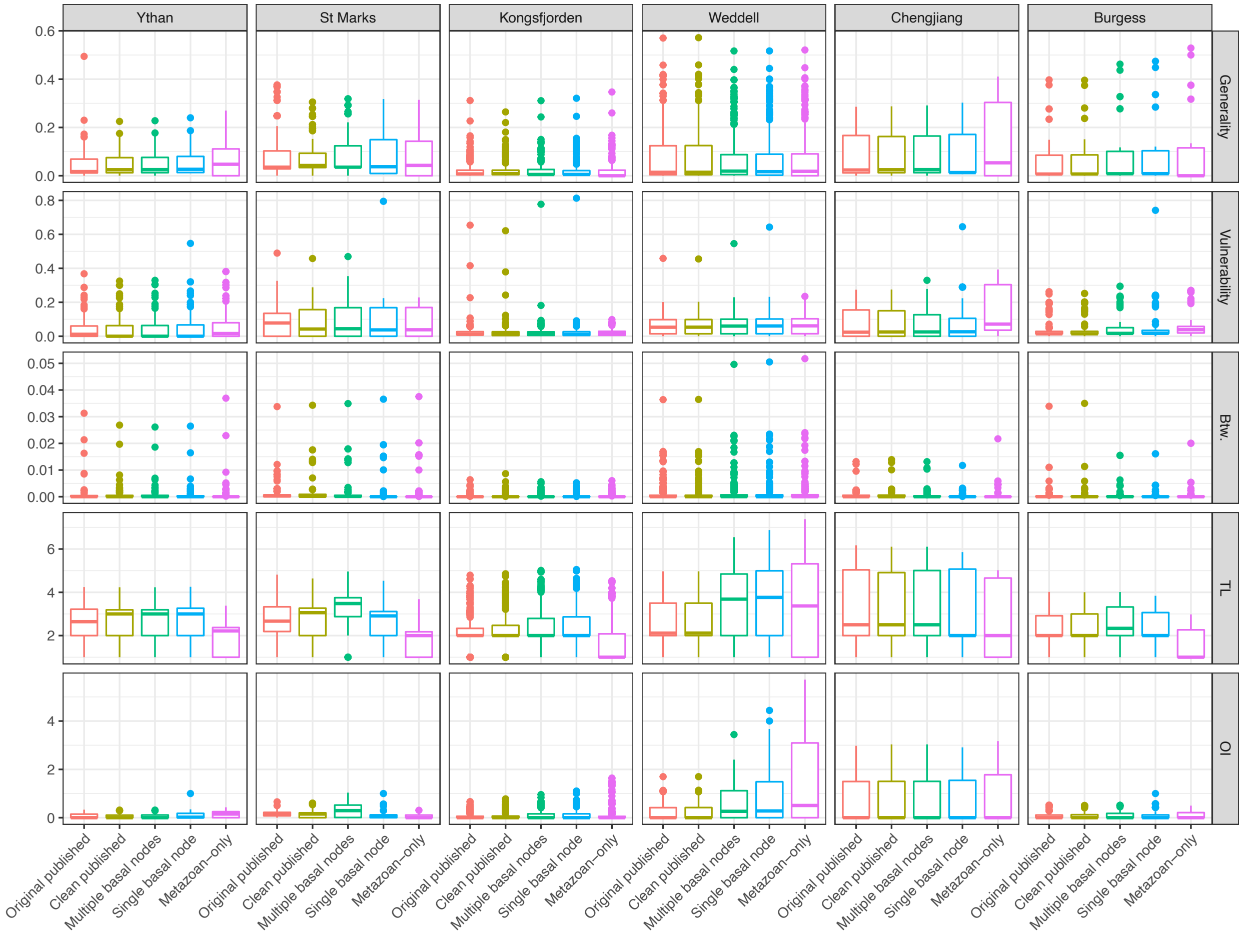

Supplementary Figure 3: Distributions of node-level metrics of various edited versions of empirical food webs considered in this study: original published webs, cleaned published webs, multiple-basal-nodes webs, single-basal-node webs, metazoan-only webs (see supplemental methods for details regarding each version type).

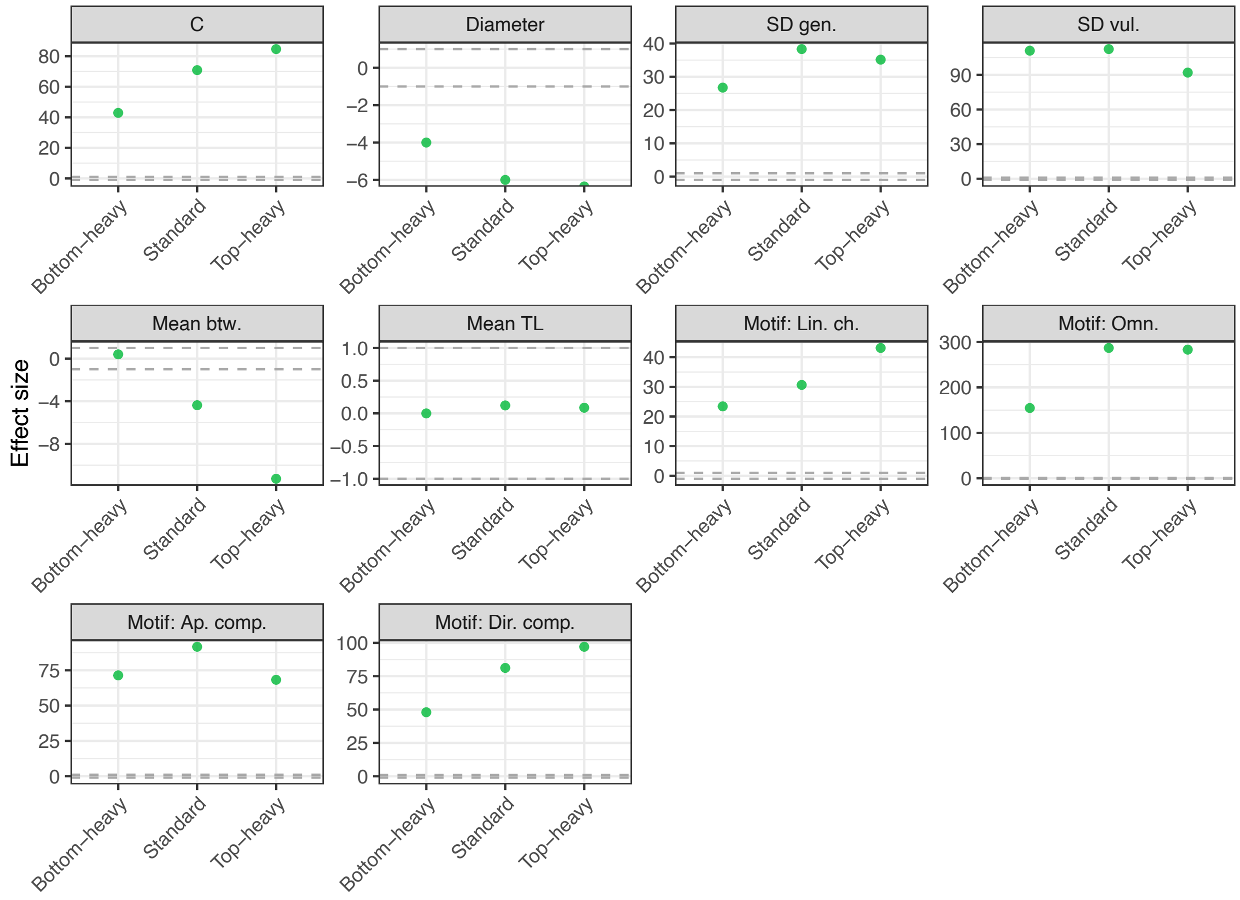

Supplementary Figure 4: Model error values describing the difference between the structure of the feasible food webs (green) and the series of replicate realized webs. Feasible web metric values were normalized based on the corresponding distribution of realized metric values.

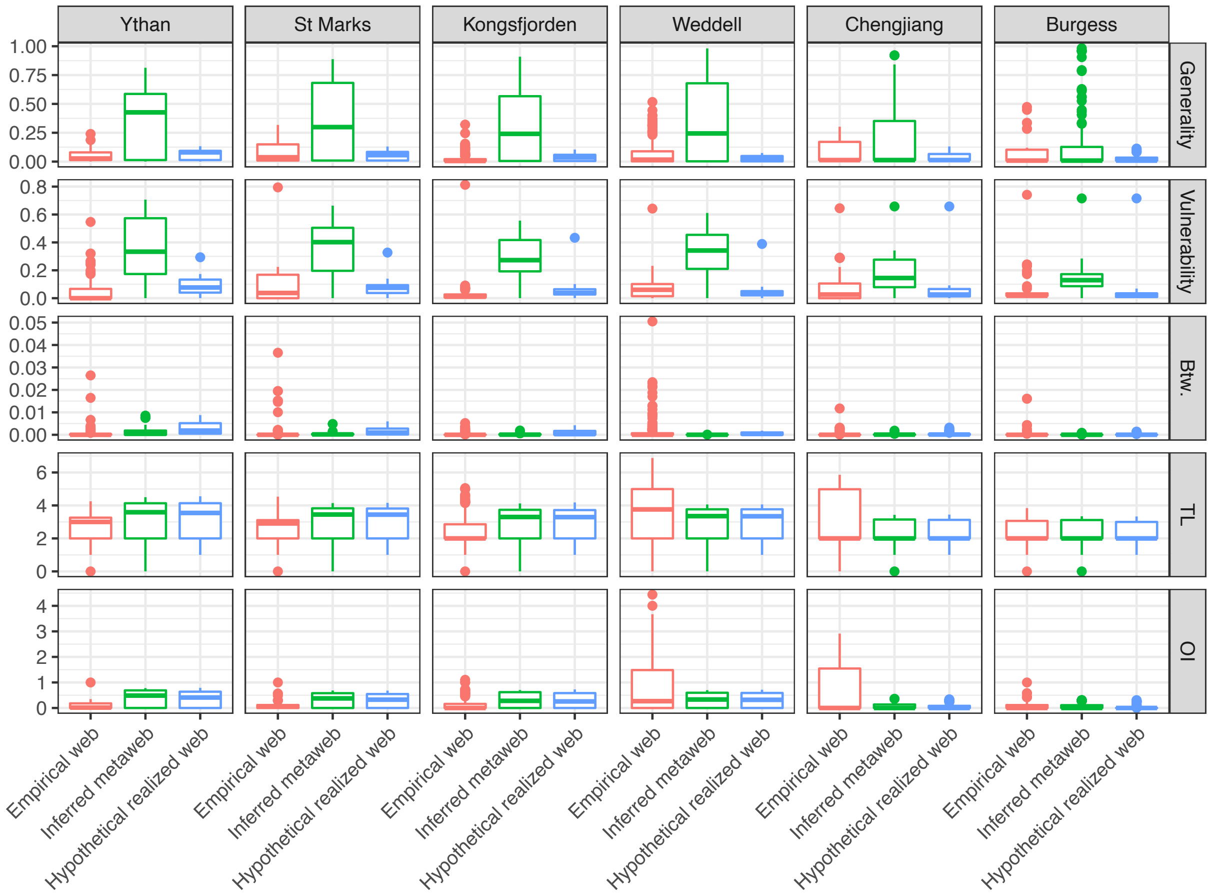

Supplementary Figure 5: Distributions of node-level metrics for empirical, feasible, and realized webs. Node-level metrics for realized webs have been averaged within-taxon for 1000 replicates (i.e., the node-level metric of Taxon A is averaged across 1000 instances of Taxon A).

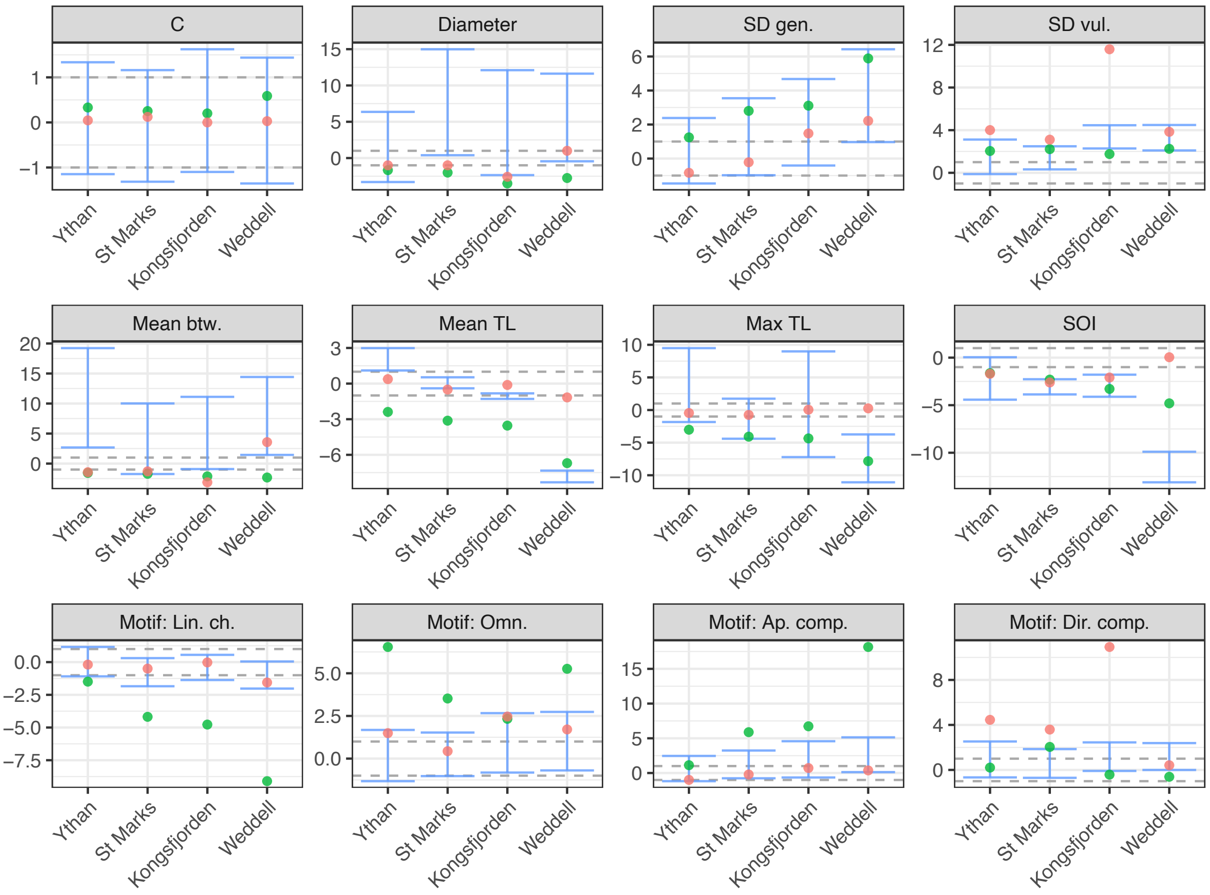

Supplementary Figure 6: Network-level statistics of niche-normalized empirical (red), feasible (green), and realized (blue) food webs. Blue error bars represent model error values of 1000 realized webs calculated as compared to 1000 corresponding niche model webs. Webs ordered by size.

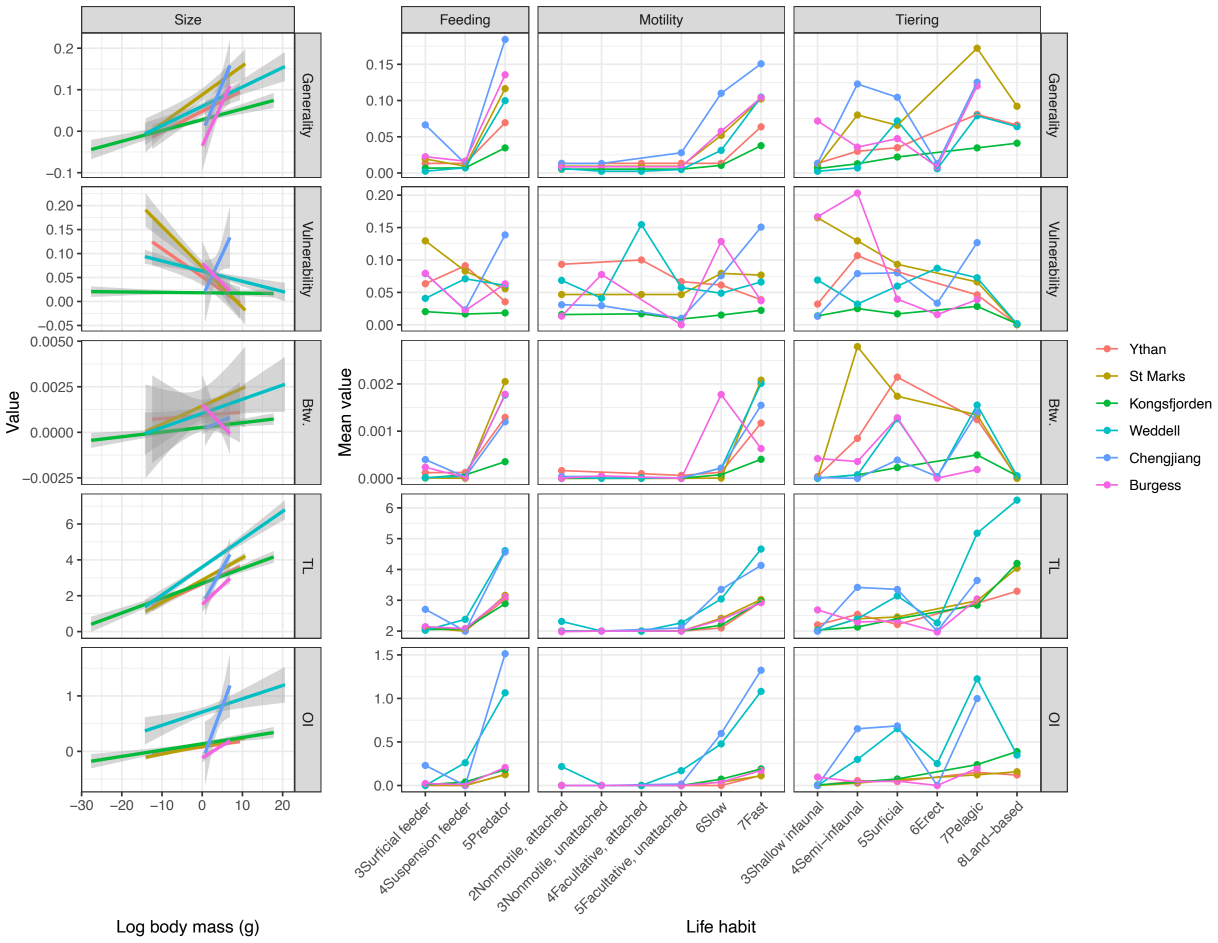

Supplementary Figure 7: Correlations between functional traits and averaged node metrics for corresponding taxa in the four focal food-webs.

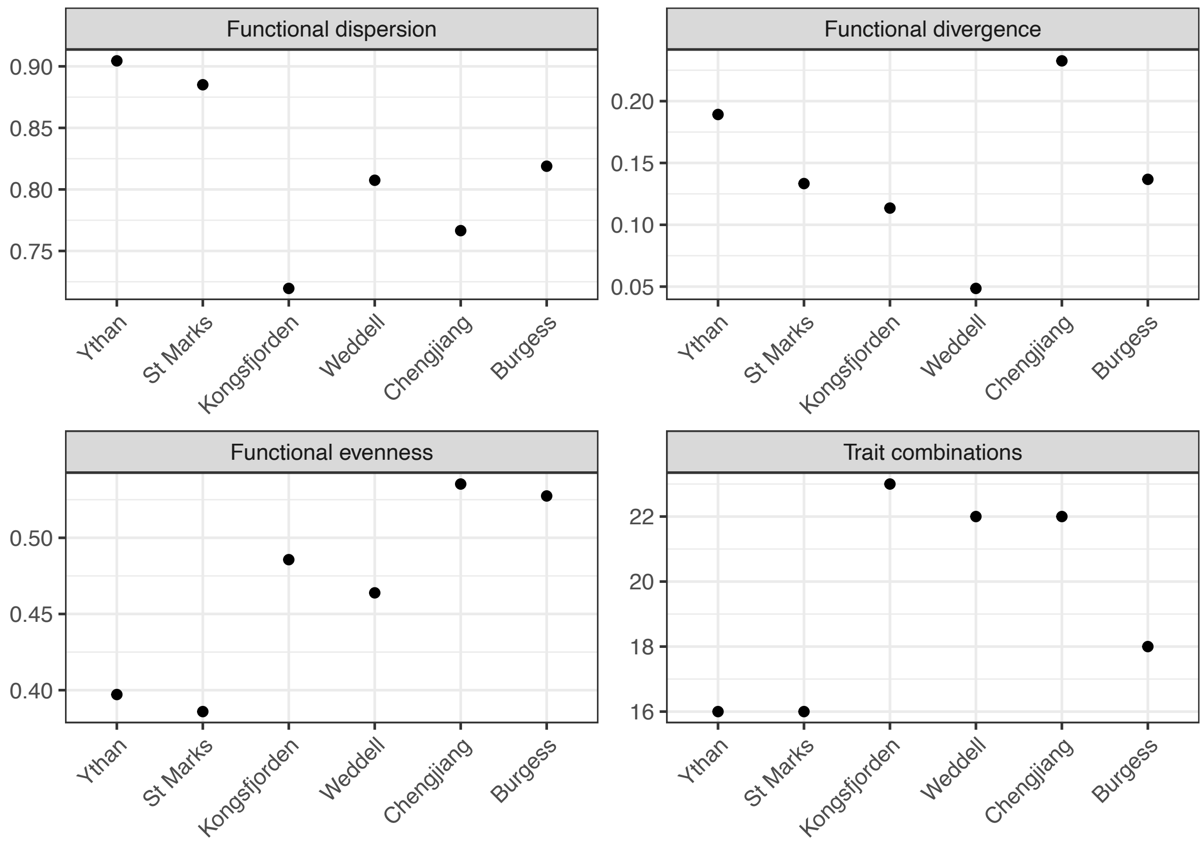

Supplementary Figure 8: Functional diversity metrics (calculated using (Novack-Gottshall 2020)) based on life habit assignments (i.e., feeding, motility, and tiering). Functional dispersion = mean pairwise distance between all taxa in niche space (i.e., PCoA axes). Functional divergence = mean distance of taxa from niche space centroid. Functional evenness = evenness of minimum-spanning-tree lengths between taxa in niche space. Trait combinations = number of unique trait combinations.

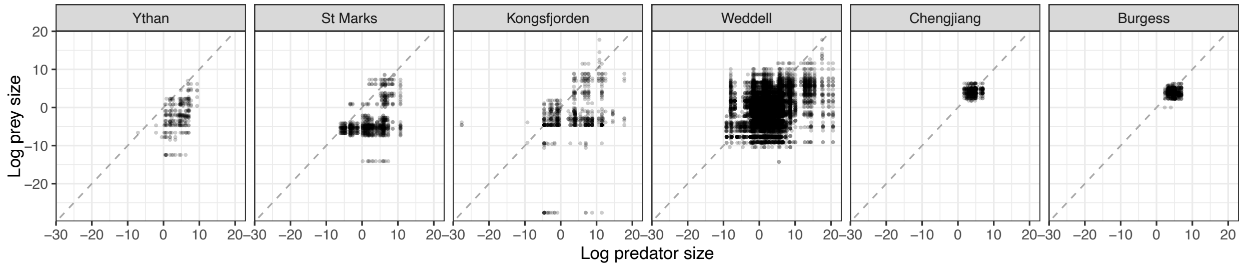

Supplementary Figure 9: Consumer-resource empirical interactions plotted by log size for the four focal food webs. Black line indicating consumer and resource sizes are equal, points above which indicate interactions where the resource taxon is larger than the consumer.

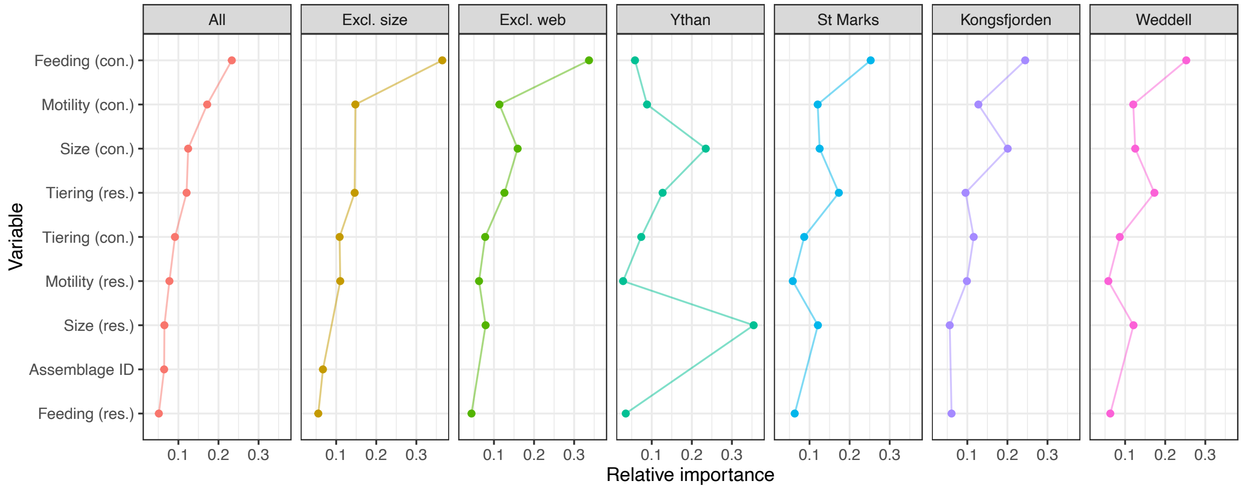

Supplementary Figure 10: Relative importance of different variables in predicting consumer-resource interactions according to a series automated machine learning models built using H2O (see supplement for more details): “All” = model built using all webs and all variables; “Excl. size” = built using all webs and all variables except for size; “Excl. web” = built using all webs and all variables except for web; web-specific models = built using each web and all variables except for web. Variables ordered by relative importance in the “all” model.

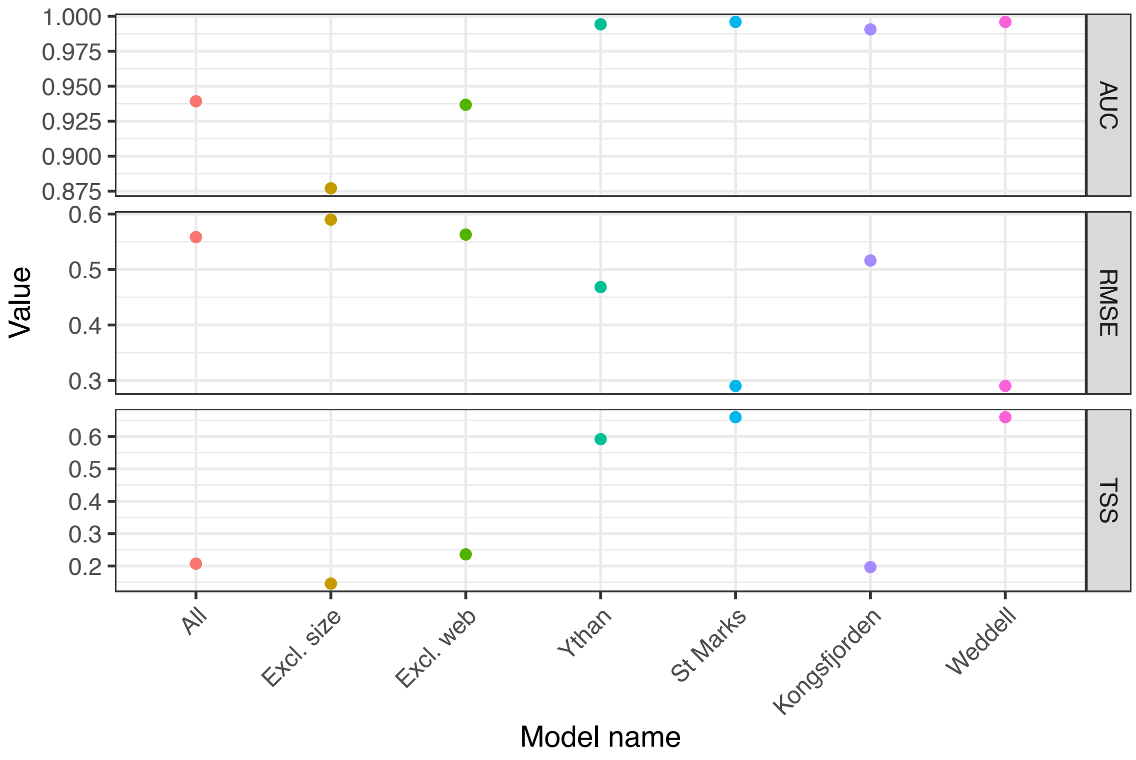

Supplementary Figure 11: Statistics for machine learning models described in Supplementary Fig. 10. “AUC” = area under curve; “r2” = r-squared value; “RMSE” = root mean square area; “TSS” = true skill statistic.

#### Supplementary tables

| Metric type | Metric | Description |
|---|---|---|
| Tiering | Semi-aquatic | Living partly in water and partly on land |
| | Pelagic | Living in the water column |
| | Erect | Benthic, extending into water column |
| | Surficial | Benthic, not extending into water column |
| | Semi-infaunal | Partly exposed to the water column, partly infaunal |
| | Infaunal | Living in the sediment |
| Feeding | Predatory | Organisms that kill individual animals that have some capacity for either protection or escape |
| | Suspension | Capturing resources from the water column |
| | Surface deposit | Capturing resources from substrate surface |
| | Primary producer | Primary producers, suspended organic matter, and detritus |
| Motility | Motile | Regularly moving |
| | Facultative, unattached | Moving when necessary, not attached to substrate |
| | Facultative, attached | Moving when necessary, attached to substrate |
| | Nonmotile, unattached | Unable to move, not attached to substrate |
| | Nonmotile, attached | Unable to move, attached to substrate |

Supplementary Table 1: Descriptions of functional traits used in this study.

#### Supplementary data descriptions

DATA WILL BE PROVIDED SEPARATELY PRIOR TO PUBLICATION

The data listed below is available at the following GitHub link:

- Food webs and corresponding taxonomic/functional trait information
- Interaction criteria to be supplied to PFIM
- R codes used for analyses in this paper
- ADBM-generated food web (see supplemental text for more information)
- List of GloBI interactions corresponding to the Ythan Estuary web (see main and supplemental texts for more information)

The PFIM package is available at the following GitHub Link:
